# Signal and Specificity of Protein Ubiquitination for Proteasomal Degradation

**DOI:** 10.1101/038737

**Authors:** Yandong Yin 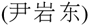, Jin Yang 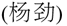

## Abstract

The eukaryotic ubiquitin system regulates essential cell events such as DNA repair, protein homeostasis, and signal transduction. Like many biochemical processes, ubiquitination must ensure signaling efficiency and in the meantime maintain substrate specificity. We examine this signal-specificity relationship by theoretical models of polyubiquitinations that tag proteins for the proteasomal degradation. Parsimonious models provide explicit formulas to key measurable quantities and offer guiding insights into the signal-specificity tradeoffs under varying structures and kinetics. Models with measured kinetics from two primary cell-cycle ligases (SCF and APC) explain mechanisms of chain initiation, elongation slowdown, chain-length dependence of E3-substrate affinity, and deubiquitinases. We find that substrate discrimination over ubiquitin transfer rates is consistently more efficient than over substrate-E3 ligase binding energy, regardless of circuit structure, parameter value, and dynamics. E3-associated substrate deubiquitination increases the discrimination over the former and in the meantime decreases the latter, further widening their difference. Both discrimination strategies might be simultaneously explored by an E3 system to effectively proofread substrates as we demonstrated by analyzing experimental data from the CD4-Vpu-SCF system. We also identify that sequential deubiquitination circuit may act as a specificity switch, by which a modest change in deubiquitination and/or processivity can greatly increase substrate discrimination without much compromise in degradation signal. This property may be utilized as a gatekeeper mechanism to direct a temporal polyubiquitination and thus degradation order of substrates with small biochemical differences.

## Introduction

A vital function of ubiquitin is to efficiently and accurately tag erroneously-synthesized, misfolded, and regulatory proteins for degradation by the 26S proteasomes [1, 2]. However, true protein substrates are usually recognized for ubiquitination via short motifs commmonly found in many nonsubstrates [3]. For example, cell-cycle ligase APC (anaphase-promoting complex or cyclosome) recognizes its substrates via D-box and Ken-box, and SCF (Skp, Cullin, F-box containing complex) via leucine rich repeats and WD40 repeats. Such low specificity in recognition motifs poses important questions: what physicochemical disparities are explored to distinguish substrates from nonsubstrates, and how the fundamental efficiency-specificity tradeoff constrains the structure and kinetics of ubiquitination circuits.

A substrate must accumulate multiple ubiquitins, possibly into a single polymeric chain or several short (sometime branched) chains, to be recognized by the 26S proteasome for degradation.This sequential ubiquitin conjugation resembles the kinetic proofreading [4, 5], a watchdog mechanism that rejects passage of nonsubstrates along a biochemical pathway. A kinetic proofreading system amplifies small biochemical disparities by introducing irreversible energy-driven intermediates, and can explain high-fidelity processes including aminoacyl-tRNA recognition by ribosome [6], antigen recognition by immune receptors [7], and signal transduction by phosphorylations [8]. Indeed, amplification of small physicochemical differences in protein ubiquitination by the collaboration of E3 ligases and deubiquitinases (DUBs) has been observed [9, 10], suggesting that some proofreading mechanisms might be encoded in the ubiquitination circuits to select substrate candidates and generate unambiguous signals. The efficiency and specificity of ubiquitination is determined by both the substrate-E3 affinity and ubiquitin transfer kinetics. The classic kinetic proofreading analysis however primarily focuses on discrimination over substrate-enzyme binding energy difference by the assumption of identical catalytic rate for both right and wrong substrates. This simple treatment is largely inadequate to protein ubiquitination. Furthermore, compared to known fidelity in DNA replication and in protein translation [11], the error rate in proteasomal degradation by protein ubiquitination remains unquantified and the relationship between ubiquitination efficiency and substrate specificity is poorly understood.

Here we develop kinetic models of varied deubiquitination schemes to examine the balance between substrate recognition and ubiquitination efficiency that underscores alternative circuit designs. Parsimoniously-parameterized models provide explicit equations to key measures including processivity, degradation signal, and specificities, which offer guiding insights into the signal-specificity tradeoffs under varying structures and kinetics. Models parameterized with experimentally-measured ubiquitination kinetics in the SCF and the APC systems quantitatively explain roles of stage-dependent ubiquitin transfer and substrate-E3 binding kinetics. We show that in all model variants proofreading over differences in ubiquitin transfer rates are substantially more rigorous than over differences in E3-substrate binding energy. DUB activity plays a pivotal role in gauging signaling efficiency and specificity. E3-associated substrate deubiquitination increases the discrimination over ubiquitin transfer and in the meantime decreases that over binding free energy, further widening their difference. Sequential deubiquitination can sharply switch on substrate specificity with little tradeoff in degradation signal, which can be utilized to establish a degradation order of substrates with small processivity differences by a common ligase.

## Results

### Model variants of different DUB schemes

Ubiquitination is catalyzed in concert by ubiquitin-activating enzyme El, ubiquitin conjugating enzyme E2, and ubiquitin ligase E3. Antagonistic to the ubiquitin conjugation, DUBs remove ubiquitins from substrates, limiting the extent of ubiquitination (Fig. 1a). We model a single substrate molecule interacting with a pool of E3 ligases. To focus on ligase-mediated substrate modification, the models collapse the cascade of ubiquitin charging to E1 (a step driven by ATP hydrolysis), ubiquitin transfer from E1 to E2, E2 binding to E3 and subsequent ubiquitin conjugation to the substrate into a single ubiquitin transfer event. Based on evidence from cell-cycle ligases, APC [9, 12] and SCF [13], we consider that a ubiquitin chain is sequentially elongated onto the substrate, one ubiquitin per transfer step. We further assume ubiquitin transfer rates are independent of downstream pathways, excluding feedback effects such as E2 degradation or ubiquitin depletion. To assess the roles of DUBs, the models examine varied DUB editing schemes: (i) *en bloc* deubiquitination (EBD, Fig. 1b), by which the DUB removes an entire polyubiquitin chain by cleaving the isopeptide bond at the substrate lysine-ubiquitin interface (endo-deubiquitination); (ii) Sequential deubiquitination (SQD, Fig. 1c), by which the DUB removes one ubiquitin at a time from the distal end of a ubiquitin chain (exo-deubiquitination). To probe the effect of E3-bound DUB editing, each model further considers whether the DUB removes ubiquitins from a substrate before it dissociates from the ligase. We anticipate that the behavior of the more general DUB scheme of removing a partial ubiquitin chain at a single step interpolates the EBD and SQD.

**FIG. 1.**
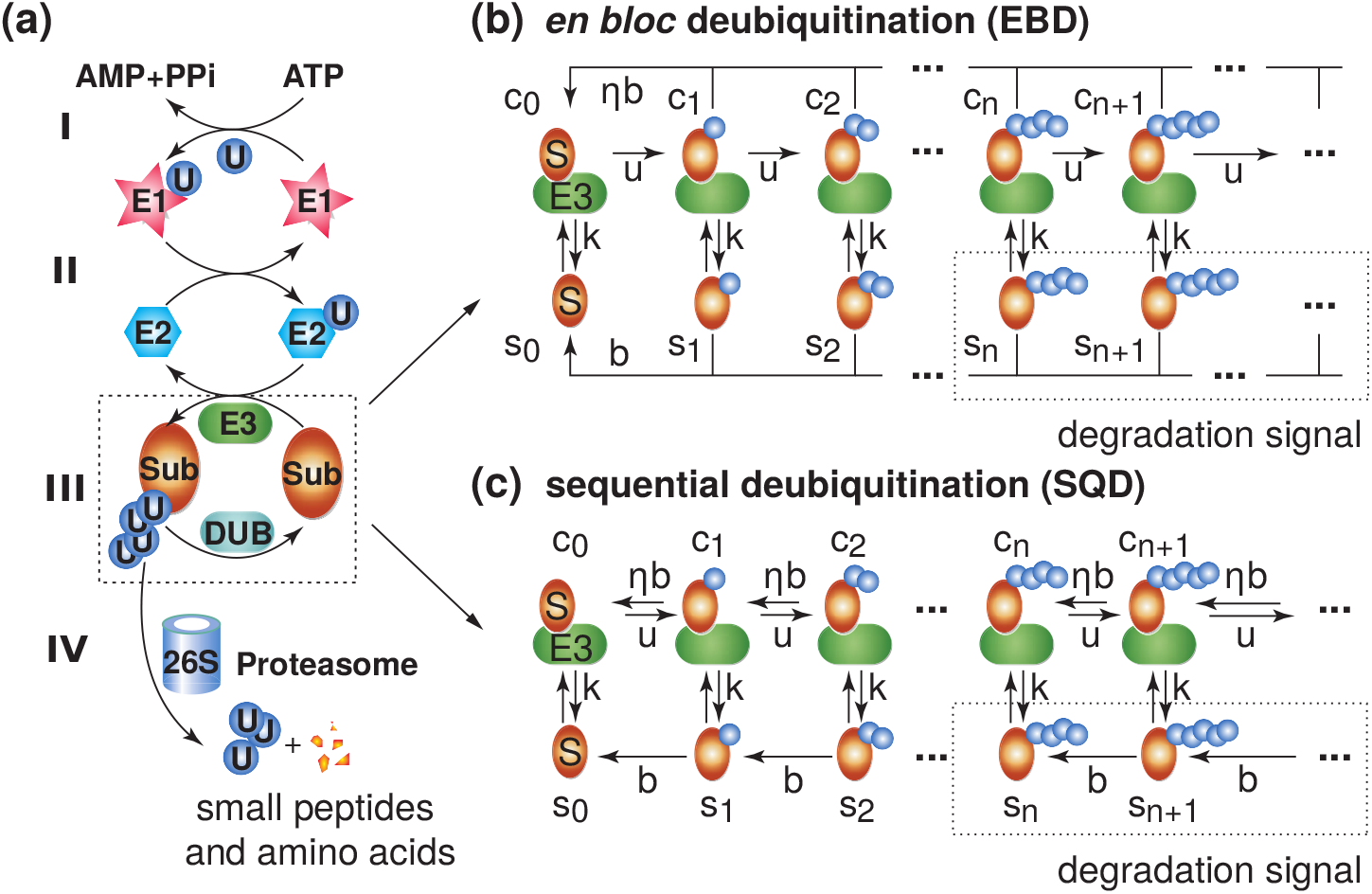
Polyubiquitination models. (a) The ubiquitin-proteasome system. At an ATP-hydrolysis step, the ubiquitin-activating enzyme E1 is charged with a ubiquitin (I), which then transfers the ubiquitin to the ubiquitin-conjugating enzyme E2 (II). The E3 ligase recognizes a substrate and bridges a ubiquitin transfer from E2 to the substrate and the succeeding chain elongation. A DUB removes ubiquitins from the modified substrate (III). Polyubiquitinated substrate is recognized by the proteasome for degradation (IV). (b) *en bloc* deubiquitination (EBD) model: A DUB cleaves the entire ubiquitin chain from the substrate by a single step. (c) Sequential deubiquitination (SQD) model: A DUB removes the ubiquitin at the distal end of the chain. *s_n_* and *c_n_* are substrate states with *n* ubiquitins dissociated from and bound to the E3 ligase and also denote the probabilities of a substrate to be in such states. A model may (*η* = 1) or may not (*η* = 0) consider E3-bound DUB editing. All rates are normalized by the 1st-order E3-substrate binding rate *k*_+_ = *k*_on_[E3], i.e., the dissociation rate constant *k* = *k*_off_/*k*_+_, the ubiquitin transfer rate *u* = *k_u_*/*k*_+_, and the deubiquitination rate *b* = *k_b_*/*k*_+_. Only uniform dissociation, ubiquitination, and deubiquitination rates are shown, which in general are ubiquitination stage-dependent.

The EBD model (Fig. 1b) resembles the Hopfield-Ninio model in the one-step backtracking, but deviates in three important aspects: (i) An intermediate state *c_n_* does not directly hop to the basal state so. The substrate first dissociates from the E3 before being reduced to basal state *s*_0_; (ii) State *C_n_* may hop to basal state *c*_0_ by E3-bound DUB editing; (iii) Dissociated substrates *s_n_* may rebind the ligase for further modification. In the absence of E3-bound DUB editing (*η* = 0), the EBD model recovers the Hopfield kinetic proofreading scheme when deubiquitination is much faster rebinding (*b* ≫ 1). The SQD (Fig. 1c) model, also without E3-bound DUB editing, resembles a ladder network used to study the dynamical instability of microtubule growth [14], where the microtubule catastrophe and rescue are analogous to dissociation and deubiquitination and to substrate-E3 rebinding and further elongation, respectively.

### Processivity, chain length, degradation signal, and specificity

Assuming stage-independent dissociation, ubiquitin transfer, and DUB rates: *k, u,* and *b*, we derive explicit formulas for most model variants (Supplementary Materials and Table S1), with the exception of the numerically-simulated SQD model under E3-bound DUB editing. The assumption of uniform rates is later relaxed when we analyse experimental data.

The *processivity* is defined as the average number of ubiquitins conjugated to a substrate during a single substrate-E3 contact. Due to the stochasticity in ubiquitin transfers and substrate dwell time per E3 contact, the number of ubiquitins conjugated to a substrate after it disengages from the ligase follows a distribution, from which the processivity and its variance can be calculated. In general, the processivity, *ρ_n_*, is a function of the initial substrate-E3 contact state *c_n_*. For the EBD model, we analytically calculate:

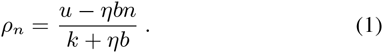

As an important and experimentally-quantifiable measure of ubiquitination efficiency, *ρ_n_* encodes influences from all kinetic processes and the ubiquitination stage *n* (Fig. 2a). High processivity is a result of fast ubiquitin transfer (large *u*) and/or long E3 ligase dwell time (small *k*). E3-bound DUB editing (*η* = 1) limits the processivity, and *ρ_n_* decreases with *n* and becomes negative when *u* < *ηbn*. Therefore, rebinding of a highly-ubiquitinated subtrate to the E3 ligase is less- or even counter-productive. Without E3-bound DUB editing (*η* = 0), both EBD and SQD models have *ρ_n_* = *ρ* = *u*/*k*, independent of the initial contact state and deubiquitination. Processivity is independent of ligase concentration and does not account for the DUB effect on dissociated substrates. For the SQD model with E3-bound DUB editing, *ρ_n_* depends on the competition between elongation and E3-bound deubiquitination (*ρ_n_* < 0 if *u* < *b* and *ρ_n_* ≥ 0 if *u* ≥ *b*), in which the substrate modification can be viewed as undergoing a one dimensional biased random walk arrested by substrate dissociation.

The *mean ubiquitin chain length* 〈*n*〉_*s*_ = Σ_*n*>0_ *ns*_*n*_/Σ_*n*≥0_ *s_n_* measures the overall ubiquitination level in dissociated substrates, whereas the *degradation signal*, *S_m_* = Σ_*n*≥*m*_ *s_n_*, depending on the minimal signaling chain length *m*, is the probability of a substrate in a state recognizable to the proteasome. We relate *S_m_* to the substrate degradation rate on the basis that proteasomal degradation involves multiple rate-limiting steps including substrate dislocation, unfolding, deubiquitination and translocation. Increasing with the protein size, degradation time at the proteasome ranges from tens of seconds to tens of minutes per protein [15, 16], whereas polyubiquitination proceeds in seconds. We obtain steady-state equations for *m* ≥ 1:

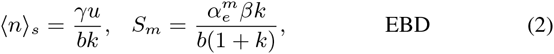

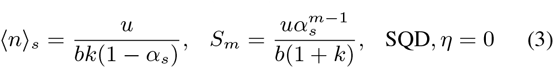

where *β* = *b*/(1+ *b*) is the probability for a dissociated and modified substrate *s_n_* to be deubiquitinated (to the ground state *s*_0_ in the EBD model, and to *s*_*n*-1_ in the SQD model). Its inverse 1/*β* is the average number of E3 contacts made by the substrate before a DUB event. *γ* = *βk*/(*βk* + *ηb*) is the probability that *c_n_* is deubiquitinated via the substrate dissociation pathway. In the EBD model *α_e_* = *u*/(*u* + *βk* + *ηb*) is the probability of an E3-bound and modified substrate *c_n_* to be incrementally modified to state *c*_*n* + 1_ without backtracking (a substrate may dissociate from the E3 and subsequently rebind). In the SQD model (*η* = 0), *α_s_* = *u*/(*β*(*u* + *k*)) is the ratio of the probability of forward walk from *c_n_* to *c*_*n*+1_ to that of backward walk from *s_n_* to *s*_*n*-1_. When *α_s_* > 1, chain elongation in the SQD model is unbounded and Eq. 3 does not apply (i.e., 〈*n*〉_*s*_ is indefinite and *S_m_* = *k*/(1 + *k*)). Substantial fluctuations 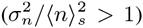 reflect high stochasticity in single-substrate ubiquitination (Eq. S5). The SQD model consistently outperforms the EBD model in generating larger degradation signal (left panel of Fig. 3a).

A ligase distinguishes modest differences in E3 binding free energy and ubiquitin transfer kinetics among its substrates. To quantify the extent of output variations caused by small energetic and kinetic differences, we define the substrate *specificity* as: Φ_*m*_ = – *∂* log *S_m_*/*∂* log *k* or Ψ_*m*_ = *∂* log *S_m_*/*∂* log *u*, which measures the degree of substrate discrimination local to a parameter value (orders of magnitude change in *S_m_* per order of magnitude change in kinetic parameter *k* or *u*). The larger Φ_*m*_ or Ψ_*m*_, the greater discrimination over difference in binding energy or transfer kinetics. This specificity measure contrasts the commonly-used ratio between turnover rates of substrate and nonsubstrate [4] and has the advantage of working with one parameter set. The EBD model has:

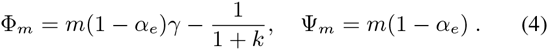

The simplicity of the equations provides essential insights. Φ_*m*_ can be interpreted as the sum of probabilities for a pre-signal state *c_n_ n* ≤ *m*, to backtrack to basal state *s*_0_ via the substrate dissociation path up to stage *m* (the first term), which is offset by signal sensitivity to the equilibrium partition of E3-bound and dissociated states determined by the substrate-ligase binding energy (the second term). By contrast, Ψ_*m*_ combines probabilities for *c_n_* to backtrack regardless of the path (either by dissociation and then deubiquitination or by direct E3-bound DUB editing). Φ_*m*_ and Ψ_*m*_ increase as *k* increases and/or as *u* decreases. Both Φ_*m*_ and Ψ_*m*_ are bounded above by *m*, which can be approached at large *k* and/or small *u* (Supplementary Materials and Fig. S2a). Overall, low processive substrates (reduced polyubiquitination) are more specific but low in signal strength, showing the signal-specificity tradeoff. Increasing the minimal signaling chain length *m* provides more presignal check points and increases specificities, again at the cost of signal.

**FIG. 2.**
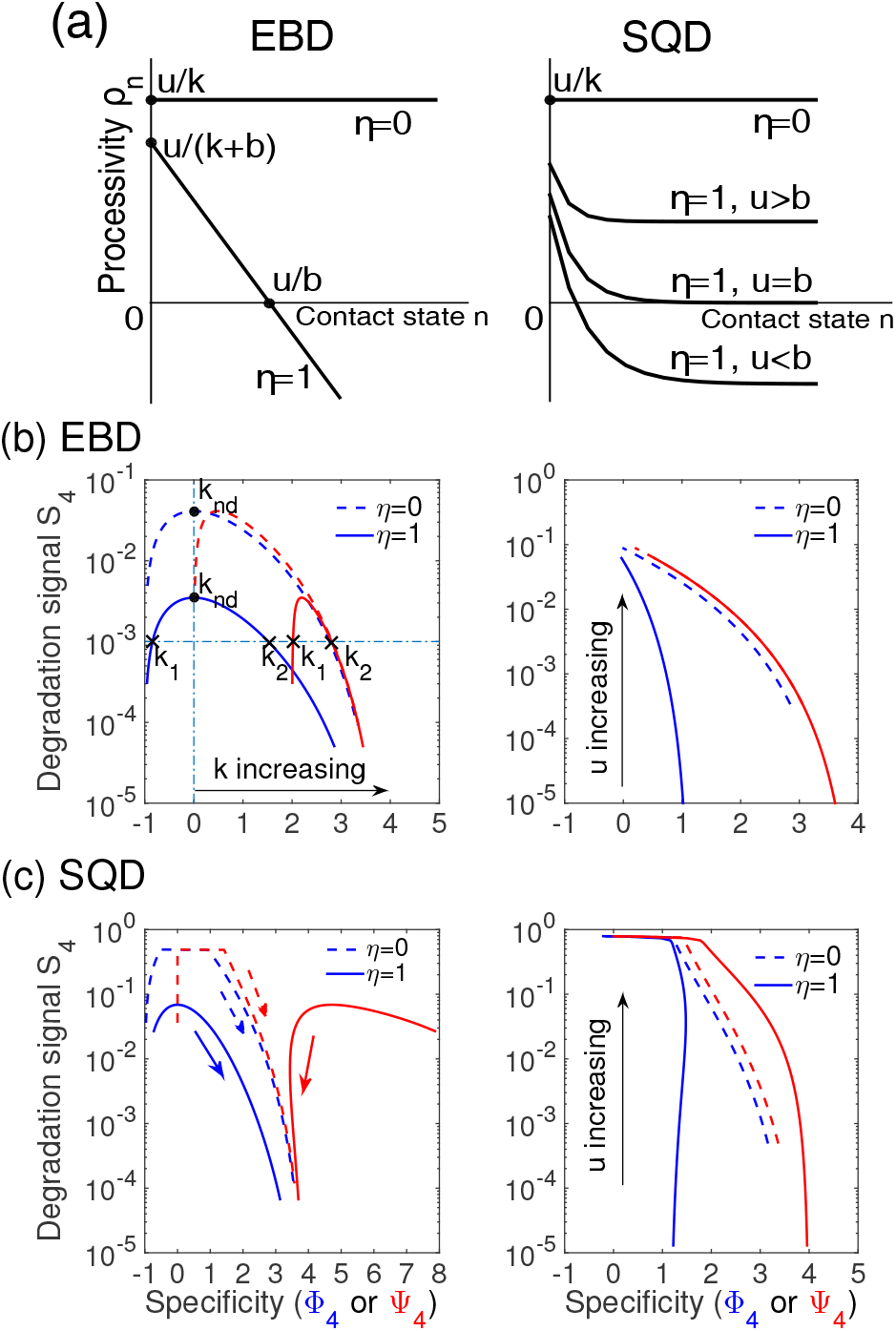
Processivity and signal-specificity tradeoffs. (a). Stage-dependent processivity in the EBD and SQD models. (b) EBD models. Left panels: *S*_4_ = 10^−3^ at *k*_1_ = 0.14 (Φ_4_ = -0.84, Ψ_4_ = 2.02) or *k*_2_ = 9.42 (Φ4 = 1.48, Ψ_4_ = 2.79) when *η* =1. The maximum signal at non-discriminative point Φ_4_ = 0 is reached at *k*_nd_ = 1.56. Right panel: *S*_4_ and Ψ_4_ has an identical tradeoff curve independent of *η* as *u* varies. (c) SQD models. Left panel: *k* increases from 0.0375 to 37.5 (arrows), and *u* = 6.25. Right panel: A maximum Φ_4_ is attained when *u* = 8.21 for the case *η* = 1. *u* increases from 1 to 62.5, and *k* = 3.75. The deubiquitination rate *b* = 6.25. Models are simulated with 400 ubiquitination stages.

The Hopfield-Ninio model proofreads against lower-affinity decoys. The EBD and SQD models however exhibit two-regime discrimination over substrate-E3 binding energy (Fig. 2b and c). In the high-affinity regime (*k* < *k_nd_*), the systems proofread against tight binding (Φ_*m*_ < 0), where the substrate is ubiquitinated with a high processivity but remains ligase-bound and inaccessible to the proteasome. In the weaker binding regime at *k* > *k*_nd_, the system discriminates against lower-affinity nonsubstrates (Φ_*m*_ > 0). Weak E3 binding reduces the processivity *ρ* and consequently decreases the fraction of degradable substrates *S_m_*. A system is non-discriminative (Φ_*m*_ = 0) at *k*_nd_ (Eq. S14), the maximum level of *S_m_* is attained. The existence of two proofreading regimes implies that two substrates of different affinity may generate identical degradation signals but differ in specificities and kinetics (Fig. 2b, left panel). This result shows that tight (lock-and-key) substrate-ligase binding should be unusual and relatively weak binding benefits substrate specificities without compromising signaling efficiency, which is supported by the existence of short E3-binding motifs of modest affinity in many substrates. A large number of tight binding substrates could also sequester the ligase population and suppress ubiquitination, which can be simulated by reducing the free ligase concentration (not shown).

**FIG. 3.**
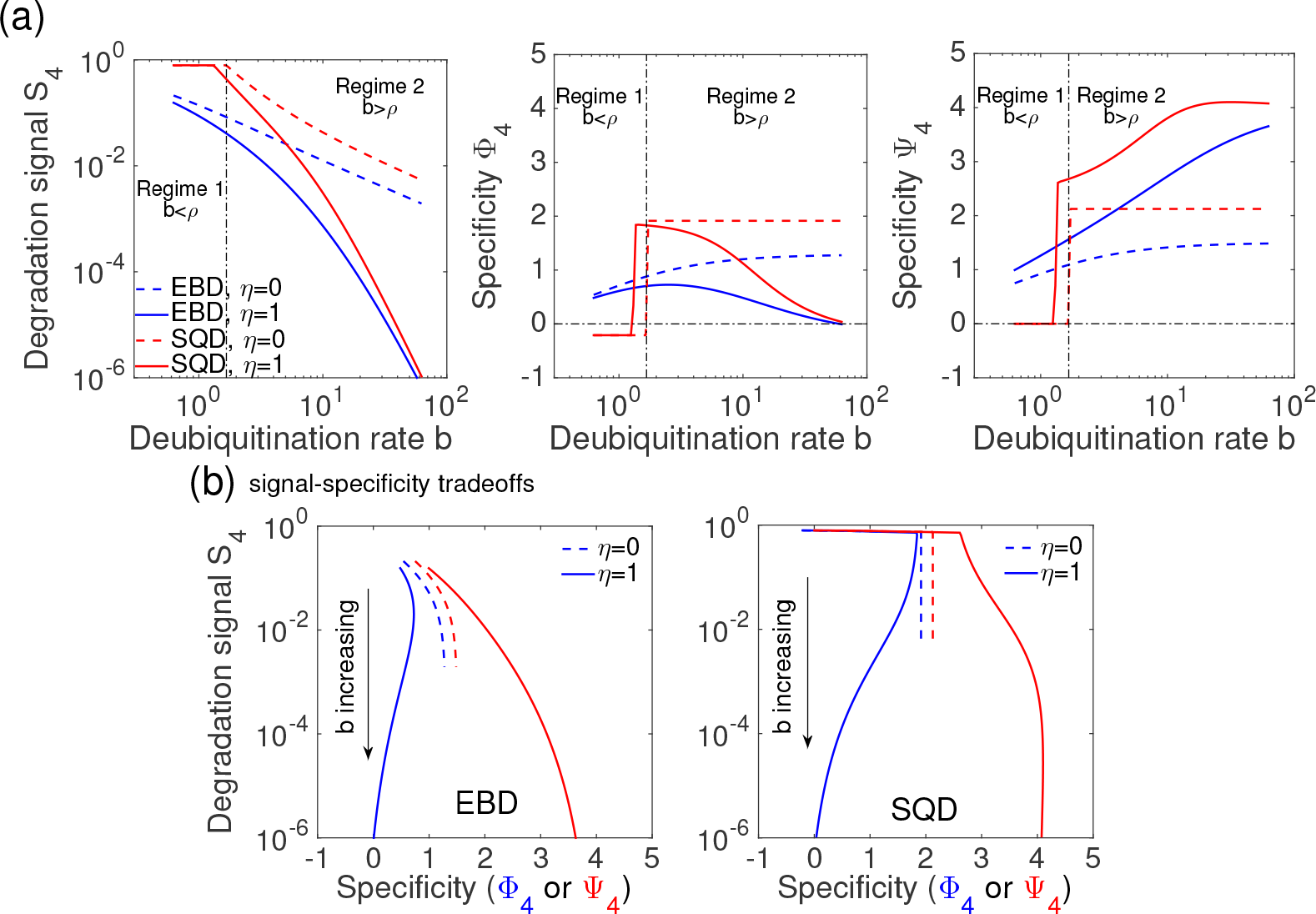
Differential EBD and SQD behaviors in the parsimonious models. (a) Degradation signal *S*_4_ vs. deubiquitination rate *b* (left panel). The SQD model (*η* = 0) is an ideal switch for specificities Φ_4_ and Ψ_4_ (middle and right panels, dashed and red), indicated by two regimes divided at the threshold *b* = *ρ* = 1.67 (dash-dotted). When *η* = 1, the DUB threshold shifts leftward. (b) Signal and specificity tradeoffs in EBD models and SQD models (b varies from 0.625 to 62.5). Other parameter values: *m* = 4, *u* = 6.25, and *k* = 3.75. Models are simulated with 400 ubiquitination stages.

An interesting property of the SQD model with E3-bound DUB editing is the existence of a discriminative regime with ultra-high specificity Ψ_*m*_. Fig. 2c shows that Ψ_4_ reaches near 8 at low *k*, which is far greater than the limit 4 in the SQD model without E3-bound DUB editing. The emergence of this high discriminative regime depends on the relative rate of elongation *u* to the DUB rate *b.* Tight substrate binding (small *k*) produces long E3 dwell time that allows the substrate undergoes a biased random walk along the elongation pathway, which generates more presignal transfer events when *u* and *b* are close (*u* = *b* = 6.25 in Fig. 2c). The strength of discrimination depends on the absolute value of *u* or *b* (Fig. S3a and b).

### Sequential deubiquitination acts as a specificity switch

For the SQD model without E3-bound DUB editing (*η* = 0), the specificities are calculated as (Supplementary Material: *SQD models – Bounded and unbounded elongation*):

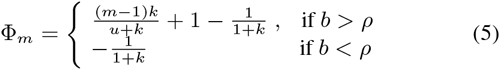

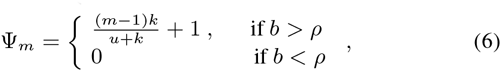

which shows that Φ_*m*_ and Ψ_*m*_ are ideally switched by DUB activity at the threshold (*b* = *ρ* = *u/k*) and are DUB-independent below (*b* < *ρ*) and above threshold (*b* > *ρ*) (Fig. 3a middle and right panels). In low DUB activity regime (*b* < *ρ* = *u/k*), the chain elongation becomes unbounded and all substrates eventually accumulate ubiquitin conjugates of size longer than the minimal length, *S_m_* + *C_m_* = 1, where *C_m_* = Σ _*n* ≥ *m*_ *c_n_.* The system proofreads against tight substrate binding with specificity Φ_*m*_ = –1/(1 + *k*), whereas the specificity Ψ_*m*_ vanishes since variations in the elongation rates in this regime do not alter the steady-state degradation signal. In high DUB activity regime (*b* > *ρ*), the chain elongation is bounded, where Φ_*m*_ and Ψ_*m*_ both switch to higher and minimal chain length- and processivity-dependent levels (Eq. 5 and Fig. 3a). The sharp switching behavior in specificity are also present in the SQD model with E3-bound DUB editing (Fig. 3a). By contrast, the specificities in the EBD model behave much gradually in response to DUB variations because the chain elongation is always bounded by one-step deubiquitination to the ground state.

### Differential E3-bound DUB editing effects on specificities

Increasing deubiquitination expectedly attenuates the degradation signal irrespective to model structures. The SQD generates higher degradation signal than EBD with or without E3-bound DUB editing at any fixed DUB activity level (Fig. 3a, left panel), and SQD mediates better signal-specificity tradeoffs than EBD. DUB effects on specificities however vary with deubiquitination schemes. Ψ_*m*_ > Φ_*m*_ universally holds at any parameter value (Eq. 4-6), signal level and DUB scheme (Fig. 2b and c. Fig. 3), showing that substrate discrimination over ubiquitin transfer rates is consistently more efficient than over binding free energy. More importantly, E3-bound DUB editing (*η* = 1) substantially widens the gaps between Φ_*m*_ and Ψ_*m*_ in all model variants. The principal circuit design idea of proofreading substrates by a biochemical process is to allow repeated passages through this process before arriving at the signaling state. A ubiquitination circuit that permits more pre-signal passages through substrate-E3 dissociation or ubiquitin transfer has a higher specificity in Φ_*m*_ or Ψ_*m*_. With E3-bound DUB editing, a modified substrate can backtrack without traversing a dissociation pathway, allowing relatively more passages of ubiquitin transfer events and therefore increasing Ψ_*m*_ at the cost of reducing Φ_*m*_. These differential DUB effects on the two specificities are verified for the EBD models by Eq. 4 or for the SQD model by simulation (Fig. 3a, middle and right panels). In the EBD model of typical parameter values: *u* = 6.26, *k* = 3.75, and *b* = 6.25, E3-bound DUB editing (*η* = 1 in Eq. 4) decreases Φ_*m*_ by 0.145m, while increases Ψ_*m*_ by 0.25m. Under the E3-bound DUB editing, an optimal DUB activity level also exists to maximize the specificity Φ_*m*_ (Fig. 3a and Eq. S17). Φ_4_ and Ψ_4_ diverge in the high DUB activity regime. The former decreases whereas the latter undergoes a second phase increase to reach a plateau, where degradation signal is inhibited (*S*_4_ < 10^−4^). High E3-bound DUB activity reduces discrimination over substrate binding energy to the minimum because deubiquitination frequently brings E3-bound substrate back to less ubiquitinated state without routing through the substrate dissociation pathway (a necessity for proofreading the difference in E3-binding energy). Forming a DUB-E3 complex potentially prevents E3 ligase from autoubiquitination, limits haphazard ubiquitinations of E2 and E3 activator proteins, or ubiquitinates the DUB itself for degradation. Our results highlight that the existence of many DUB-E3 association pairs [17] might be purposed for proofreading ubiquitin transfer kinetics, which as we demonstrated is far more efficient than proofreading binding energy.

**FIG. 4.**
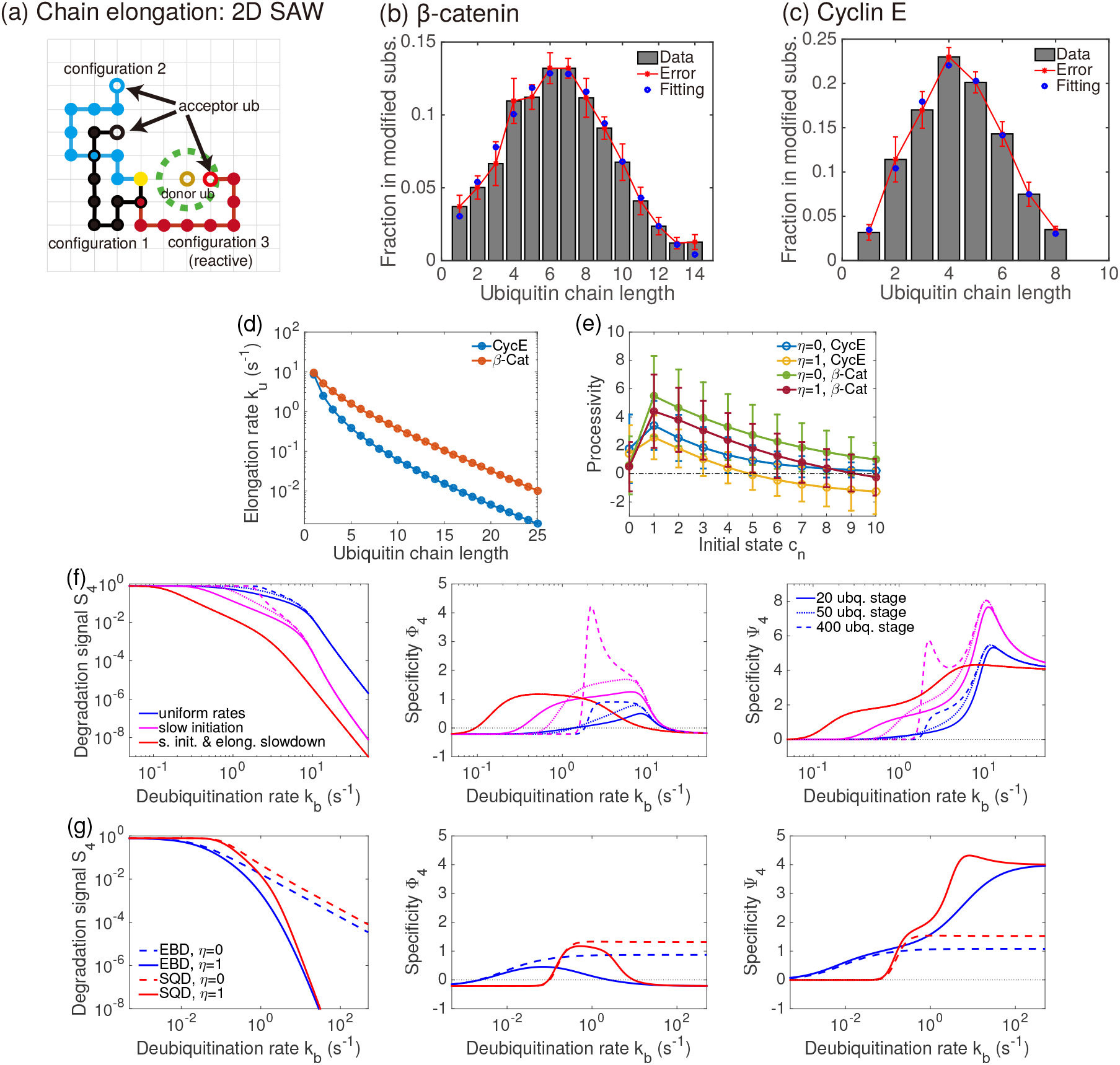
Ubiquitin chain initiation and elongation. (a) A 2D lattice illustration of the SAW model of chain elongation. Three possible configurations of a ubiquitin chain (*n* = 9). The acceptor ubiquitin (red) in configuration 3 (green) is inside the reactive volume (dashed circle) around the donor ubiquitin. (b) and (c) Fits to single-contact chain length distributions (excluding the unmodified substrates) of the SCF substrates [13]: *β*-catenin by Cdc34-SCF^*β*-TrCP^ (initiation rate *k*_*u*,0_ = 0.03 s^-1^ and elongation rates *k_u,n_* = 11.74 x 1.223^-*n*^ x *n*^-0.627^ s^-1^), and Cyclin E1 by Cdc34-SCF^Cdc4^ (*k*_*u*,0_ = 0.2 s^-1^ and *k_u,n_* = 9.78 x 1.166^-*n*^ x *n*^-1.58^ s^-1^). (d) Elongation rates. See Supplementary Materials: *Ubiquitination models – SAW model of elongation slowdown* for the fitting procedure. (e) The stage-dependent processivity for cyclin E and *β*-catenin. ODE SQD models (*k_b_* = 0.5 s^-1^) with all *s_n_* as absorbing states are solved to the steady state with initial conditions *c_n_* = 1, for *n* = 0, 1,10. Processivity from each starting state and its variance are then calculated as 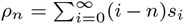 and 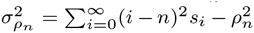 (s.e.m. shown as error bars). (f) Signal-specificity tradeoffs (left) for the parsimonious model (blue, all ubiquitin transfer rates equal to *k*_*u*,1_), slow initiation model (magenta, chain initiation rate *k*_*u*,0_ = 0.03 s^-1^ and others equal to *k*_*u*,1_ of *β*-catenin), and slow initiation and elongation slowdown (red, elongation rates *k*_*u,n*_). Specificity Φ_4_ (middle) and Ψ_4_ (right) vs. *k_b_*. Steady-state probability distribution was solved by a linear system of a finite-sized SQD model with 20 (solid), 50 (dotted), or 400 (dashed) ubiquitination stages to compute the degradation signal and specificities (Supplementary Materials: *Numerical computation of model quantities*). Simulation results from models without elongation slowdown are more sensitive to the system size and have sharper responses to DUB activity changes when the infinite ubiquitination steps are approximated by a large finite system (400 ubiquitination stages, dashed curves.). (g) *S*_4_ (left), Φ_4_ (middle), and Ψ_4_ (right) vs. *k_b_* in EBD (blue) and SQD (red) models. Ubiquitin transfer rates are based on SCF substrate *β*-catenin (50 ubiquitination stages used in simulation). Common parameters: E3-substrate association and dissociation rate constants: *k*_on_ = 0.01 nM^-1^s^-1^ and *k*_off_ = 0.3 s^-1^, and the free ligase concentration [E3] = 8 nM.

### Chain-length dependence of ubiquitin transfer

Although the parsimonious models are efficient in offering analytical insights, kinetics of *in vitro* ubiquitin conjugation is elaborate. The first ubiquitin conjugation (chain initiation), in some cases mediated by specialized E2, is 1 to 2 orders of magnitude slower than the subsequent elongation that progressively slowdowns as the substrate accumulated more ubiquitins [12, 13, 18–21]. In a coarse-grained model, we consider the chain elongation as a self-avoiding walk (SAW) process that treats the spatial fluctuation of a ubiquitin chain as a search by the distal acceptor ubiquitin for the E2-bound donor (Fig. 4a). Under this interpretation, the elongation rate is physically limited by the chain length and is inversely proportional to the total search volume of the chain characterized by *k_u,n_* ∼ *μ*^-*n*^*n*^-*a*^ (Supplementary Materials: *SAW model of elongation slowdown*). To extrapolate the elongation rates in a specific system, we use this model to fit the bell-shaped chain length distributions (compared to the geometric distributions in the parsimonious models) measured by single-contact assays for *β*-catenin (Fig. 4b) and cyclin E (Fig. 4c) in Ref. [13]. The elongation rates decline rapidly with the chain length (Fig. 4d, a more than 5 fold decrease from *k*_*u*,1_ = 11.4 s^-1^ to *k*_*u*,4_ = 2 s^-1^ for cyclin E). The processivity from state *c*_0_ is much smaller than that from *c*_1_ due to the slow initiation (*ρ*_0_ = 0.5 vs. *ρ*_1_ = 4.5, for *β*-catenin) and progressively decays as the elongation slows down (Fig. 4e). *ρ_n_* reaches below zero when the E3-bound DUB rate exceeds the ubiquitin transfer rate (for cyclin E, *ρ_n_* < 0 when *n* > 5).

A slow chain initiation imposes a predominant kinetic barrier to filter substrates, as low conjugation rate enhances discrimination over both the binding free energy and ubiquitin transfer. Rapid elongation that efficiently generates a polyubiquitin conjugate and in the meantime may provide additional proofreading. Indeed, theoretical analysis of the EBD and SQD models (Supplementary Materials: *Chain initiation*) shows that slow initiation increases specificities Φ_*m*_ and Ψ_*m*_ at the expense of degradation signal *S_m_.* To illustrate combined effects of slow chain initiation and elongation slowdown, we simulate and compare SQD models with E3-bound DUB editing of three different kinetics under varying DUB activity: (i) uniform ubiquitin transfer rates, (ii) slow chain initiation with uniform elongation rates, and (iii) slow chain initiation with elongation slowdown. In practice a model with a finite number of ubiquitin steps is computed to obtain the steady-state results and the choice of system size may influence the outcome and therefore the interpretation of results (Fig.4f). The degradation signal expectedly decreases as the DUB rate increases and is further attenuated by slow chain initiation and elongation slowdown (Fig.4f, left panel), which shifted the specificities vs. *k_b_* relationship to a lower and physiological DUB regime (Fig.4f middle and right panels). Consistent with the parsimonious models, the specificity of ubiquitin transfer Ψ_4_ is greater than that of E3-substrate binding Φ_4_ across the entire range of DUB activity. At low DUB activity, both specificities are at their minima as substrates are indiscriminately ubiquitinated to the signaling states. With slow initiation and elongation slowdown, increasing DUB activity elevates both specificities to plateaus where they are less sensitive to DUB variations. In this DUB regime, we predict that the DUB level should be regulated near the lower bound (*k_b_* = 0. 2 – 0.3 s^-1^) to allow efficient ubiquitination (*S*_4_ ≈ 0.1). The trends of Φ_*m*_ and Ψ_*m*_ diverge at the high DUB regime (*k_b_* > 1 s^-1^), where Φ_4_ declines while Ψ_4_ increases as DUB activity increases further. Furthermore, the DUB-toggled specificity switches are preserved in the SQD model under slow initiation and elongation slowdown (with or without E3-bound DUB editing), compared to the graded responses seen in the EBD model (Fig.4g, middle and right panels). Above the DUB threshold (about *k_b_* = 0.5 s^-1^), the SQD model outperforms the EBD counterpart in both degradation signal and specificities and provides better signal-specificity tradeoffs (Fig.4g).

**FIG. 5.**
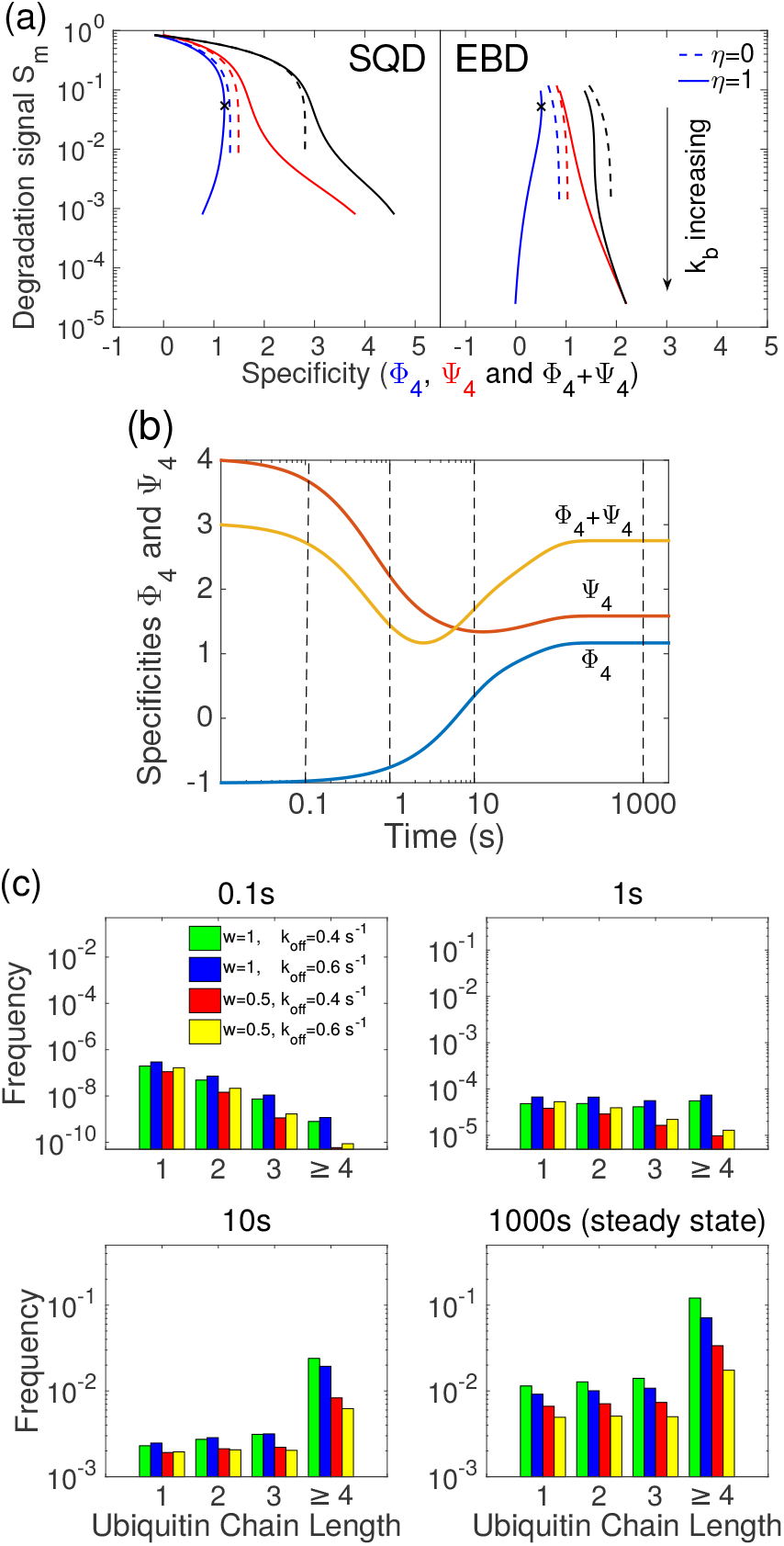
Effects of DUBs on ubiquitination efficiency and specificity. (a) DUB-regulated degradation signal-specificity tradeoffs in the CD4-Vpu-SCF system. Ubiquitin transfer rates (*k*_*u*,0_ = 0.05 s^-1^, and *k_u,n_* = 13.68 x 1.238^-*n*^ x *n*^-0.157^ s^-1^, *n* ≥ 1) fit to those adopted in the model in ref. [10] (Fig. S4). *k_b_* varies from 0.04 to 4 s^-1^ at *k*_off_ = 0.4 s^-1^. With E3-bound DUB editing (*η* = 1), the maximum specificity over binding affinity (marked by x) 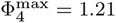 at *k_b_* = 0.79 s^-1^ (SQD) or 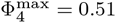 at *k_b_* = 0.087 s^-1^ (EBD). The additive Φ_4_ + Ψ_4_ is also shown to illustrate the combined effect by Φ_4_ and Ψ_4_. (b) Time trajectories of Φ_4_, Ψ_4_, and Φ_4_ + Ψ_4_, at *k_b_* = 0.5 s^-1^. (c) Polyubiquitination of four substrates: *k*_off_ = 0.4 s^-1^ or 0.6 s^-1^, and ubiquitin transfer rates *k*_*u,i*_ (proportional factor *w* = 1) or a proportional 50% (*w* = 0.5) decrease in *k_u,i_.* Chain length distributions are shown at time 0.1 s, 1 s, 10 s and 1000 s. (b) and (c) were results from the SQD model with E3-bound DUB editing. Models were simulated with the initial condition *s*_0_ = 1 and 0 to all other states (with 101 ubiquitination steps). [E3] = 8 nM and *k*_on_ = 0.01 nM^-1^s^-1^.

### Combined discrimination over ubiquitin transfer and E3-substrate binding – analysis of experimental data

A recent study of endoplasmic reticulum associated degradation by Zhang *et al.* [10] demonstrated that deubiquitination amplified small biochemical difference between wild-type CD4 and its transmembrane-domain (TMD) mutant (CD4-M). The latter has a modest 30% reduced affinity to the HIV protein (Vpu) that in its phospho-form bridges the substrate to the E3 ligase SCF for ubiquitination onto the CD4 cytosolic tail (similar results were also observed between the wild-type CD4 and Vpu TMD mutant pair). In the absence of DUBs, wild-type and mutant CD4 were almost indistinguishably ubiquitinated in vitro, especially in short-chain conjugates (*n* < 10), therefore producing near identical degradation signal. Differences in long-chain conjugates were detected due to accumulated processivity differences in wild-type and mutant CD4, which however could not isolate the contribution by substrate affinity difference from difference in ubiquitin transfer rates. By contrast, in cultured cells CD4-M ubiquitination was 5 to 10 fold less than wild-type CD4 and its degradation was nearly abolished (by a 90% reduction from the wild-type CD4). The study attributed this amplified difference to that DUBs cleave ubiquitins from low-affinity CD4-M due to its frequent dissociation from the phospho-Vpu-SCF ligase complex. Quantitative analysis by our model suggests that difference in substrate-E3 binding energy alone cannot sufficiently explain the observed difference in substrate polyubiquitination across a wide range of DUB concentration. To fully account for the data, the model predicts that the TMD mutation in CD4 may have also attenuated ubiquitin transfer rates and thus reduced the processivity to attain the multi-fold reduction in CD4-M polyubiquitination.

In agreement with the theory derived from the parsimonious models, simulation results of the SQD and EBD models with parameter values adopted from Ref. [10] (Fig. S4) showed that specificity Ψ_4_ is consistently larger than Φ_4_ across the whole DUB range. E3-bound DUB editing further amplifies the difference between Φ_4_ and Ψ_4_ (Fig. 5a). The maximum specificity 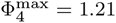 in the SQD model with E3-bound DUB editing (the same model structure used in Ref. [10]), which implies that a mutant with half the E3 affinity (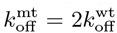) is polyubiquitinated S4 about 40% of or 2.5-fold less than the wild type, substantially inadequate to explain the observed 5-10 fold difference. Φ_4_ is even lower in the EBD models at any level of degradation signal (Fig. 5b). Explaining this degree of substrate discrimination solely by the amplification of the modest difference in binding energy (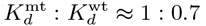) reported in Ref. [10] requires at least Φ_4_ = 4.51, which exceeds the upper limit set by the minimal signaling chain length 4. Realistic specificities are typical far below their upper limits within a broad range of DUB activity (Fig. 5a) and ubiquitin transfer rates (Fig. S5) because the signal-specificity tradeoffs prevent the system from attaining high specificities without a substantial reduction in degradation rate. This calculation suggests that specificity Φ_4_ could only partially account for the polyubiquitination difference between CD4-M and the wild type CD4, and that the rest of the discrepancy might be attributed to some other mechanism(s). One primary model prediction is that aside from weakening the affinity to Vpu, TMD mutation in CD4 might likely have reduced the efficiency of chain initiation and/or elongation, which is amplified by the proofreading. The two specificities Φ_4_ and Ψ_4_ might be both responsible for the substrate discrimination. With E3-bound DUB editing, proofreading the sequential ubiquitin transfer provides a more efficient discriminative mechanism (Fig. 5a, Ψ_4_ = 1.59 vs. Φ_4_ = 1.17 at *k_b_* = 0.5 s^-1^). Given the affinity change in CD4-M and 5-10 fold change in polyubiquitination, we can estimate that the ubiquitin transfer rates of CD4-M are about 31-47% of the wildtype ubiquitin transfer rates by the SQD model with E3-bound DUB editing (Eq. S15). Although we may not anticipate this prediction to be precisely quantitative, the difference in ubiquitin transfer kinetics between wild-type CD4 and its TMD mutant can be experimentally probed by measuring and comparing the stage-dependent processivities in the wild-type CD4 and CD4-M using a single substrate-E3 encounter assay given known *k*_off_’s of CD4 and CD4-M.

In the SQD model with E3-bound DUB editing, high DUB activity decreases Φ_4_ from 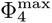 when *k_b_* > 0.66 s^-1^ and thus compromises substrate discrimination over binding affinity. By contrast, Ψ_4_ increases monotonically with the DUB rate, which compensates the loss in Φ_4_ as the additive specificity Φ_4_ + Ψ_4_ also increases with *k_b_* (Fig. 5a). DUB dosage-response data [10] did show a monotonic action of DUB, in which higher DUB concentration renders greater discrimination against CD4-M. We note, in this regime, the DUB rate is higher than reported *k*_cat_ = 0.35 s^-1^ [22] and *k*_cat_ = 0.53 s^-1^ [23] for UPS2-CD used in Ref. [10]. Asa result, the system may behave at the regime below the optimal *k_b_*, where increasing DUB concentration induces stronger substrate discrimination over binding energy. An alternative explanation is that the CD4-Vpu-SCF system has a non-significant E3-bound DUB activity, for which the monotonicity between DUB concentration and substrate discrimination can be accounted for by the model without E3-bound DUB editing (Fig. 5a).

Besides the steady state, transient behavior provides more information to ubiquitination dynamics (Supplementary Materials: *Numerical computation of model quantities*). We simulate the time evolution of specificities in the SQD model with E3-bound DUB editing (Fig. 5b). Φ_4_ changes from -1 at the start (discrimination against stronger binding because tight binding has a low *k*_off_ and thus has slow initial kinetics in generating degradation signal) to 1.17 at the steady state (discrimination against weaker binding). On the opposite, Ψ_4_ downslides from its upper limit 4 at the start to the steady-state 1.59. Such differential dynamics of the two specificities indicates the change of proofreading preference over time and raises an interesting perspective. Strong discrimination over ubiquitin transfer kinetics can be achieved at a far more rapid proteasomal degradation, or by a mechanism (dislocation or sequestration) that quickly arrests the polyubiquitinated substrate from the ligase system. Substantial discrimination against high binding energy achieves near the steady state and thus at a low proteasomal degradation rate, where combined proofreading over dissociation and ubiquitin transfer (Φ_4_ + Ψ_4_) renders more rigorous discrimination. Ψ_4_ is always higher than Φ_4_ during the transient phase. To illustrate polyubiquitination differences among distinct substrates, Fig. 5(c) shows snapshots of chain length distributions of four substrates: (1) the substrate used in Fig. 5a, (2) a substrate with 50% increase in *k*_off_, (3) a substrate with 50% decrease in *k_u,n_*, and (4) a substrate with 50% increase in *k*_off_ and 50% decrease in *k_u,n_.* At very short time of 0.1 s and 1 s, substrate 2 with a weaker affinity and a faster kinetics accumulates the highest degradation signal, while those of substrate 3 and 4 are strongly suppressed. Signal of substrate 1 exceeds that of other substrates as the system relaxes to the steady state. At the steady state, substrates 3 and 4 reach higher in relative degradation strength as they are progressively less discriminated over their ubiquitin transfer rates (see Table S2 for steady-state values).

**FIG. 6.**
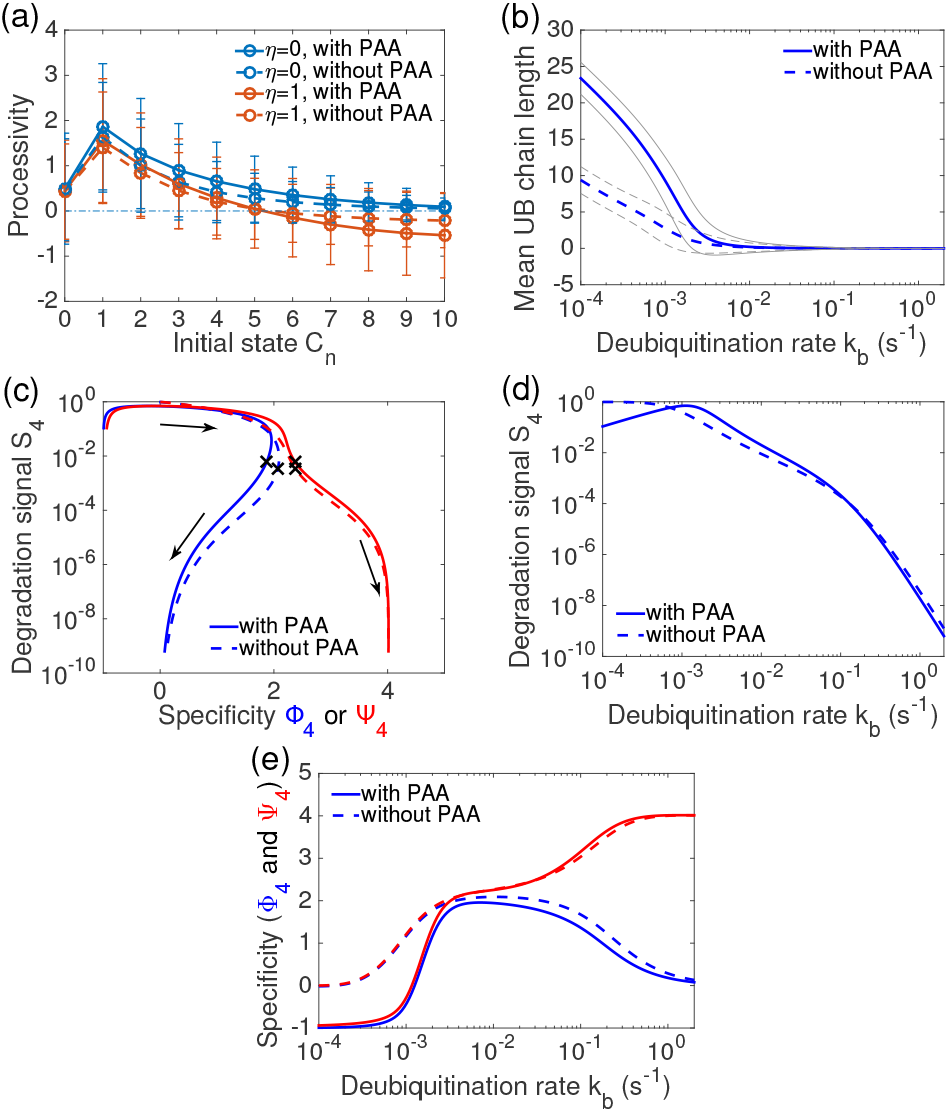
Effects of step-dependent substrate-E3 affinity on processivity, degradation signal, and specificities. (a) Processivity of APC substrate securin in Ref. [3] in the absence (dashed) or presence (solid) of PAA, with (*η* = 1) or without (*η* = 0) E3-bound DUB editing. Error bars indicate the s.e.m. *k_b_* = 0:02 s^-1^. (b) The mean chain length 〈*n*〉_*s*_ (blue) enclosed by standard deviation (grey outlines), with or without PAA. (c) Signal and specificity tradeoffs under varying DUB activity (arrows point to increasing *k_b_*. Markers ‘×’ locate at *k_b_* = 0.02 s^-1^). Φ_4_ and Ψ_4_ were computed as the relative sensitivities of *S*_4_ to proportional factors of *k*_off,*n*_ and *k_u,n_*. See Fig. S8 for fitting functions to *k*_off,*n*_, *k*_on,*n*_, and *k_u,n_*. (d) Degradation signal and (e) specificities as functions of *k_b_*. *k_b_* increases from 10^−4^ s^-1^ to 2 s^-1^. Models were simulated with 100 ubiquitination steps with [E3] = 10 nM.

### Chain-length dependence of substrate-E3 affinity

A recent single-molecule fluorescence study of APC-mediated ubiquitination by Lu *et al*. [3] observed a mechanism, namely processive affinity amplification (PAA), in which substrate affinity to APC progressively increases with the number of conjugated ubiquitins (spanning about 1 order of magnitude change in affinity among substrates with ubiquitin conjugates of varied size), by decreasing *k*_off_ (from *k*_off,0_ = 0.07 s^-1^ to *k*_off,15_ = 0.005 s^-1^) and meanwhile increasing *k*_on_ (from *k*_on,0_ = 1.3 x 10^5^ M^-1^s^-1^ to *k*_on,15_ = 4.9 x 10^5^ M^-1^s^-1^) (see Fig. S8 for fits to data from Ref. [3]). Presumably due to increased multiplicity in the ubiquitin binding interface to the ligase, the PAA allows more frequent substrate rebindings to the E3 and longer E3 dwell times, by which a modified substrate further acquires ubiquitins more efficiently to counterbalance the DUB activity. To analyze the quantitative significance of PAA, we simulate the SQD models to examine how the PAA affects the degradation signal and substrate specificities.

Similar to SCF substrates (Fig. 4e), the stage-dependent processivity of securin markedly increases after chain initiation and then declines thereafter because of the elongation slowdown (Fig.6a). Without E3-bound DUB editing, the processivity expectedly remains positive, where the PAA offers the substrate a longer ligase dwell time and therefore a higher processivity at each stage. By contrast, under E3-bound DUB editing, the PAA increases higher backward processivity for highly-modified substrates because the elongation slowdown reduces the forward processivity and the extended ligase dwell time exposes the substrate more opportunities for deubiquitination. This indicates that the PAA does not improve ubiquitination efficiency beyond a certain chain length (*n* > 5 in Fig. 6a for securin). As ubiquitination, deubiquitination, and dissociation are random processes, the stochasticity in processivity and low DUB activity further obscures the difference caused by the PAA. At low DUB activity, even though the PAA produces extended elongation reflected in the mean chain length in dissociated substrates (〈*n*〉_*s*_, Fig. 6b), it generates lower degradation signal than the system without PAA due to tight binding on the ligase (Fig. 6d). The PAA modestly improves polyubiquitination signal *S*_4_ at a slight cost of specificity Φ_4_ (Fig. 6c, d and e), consistent with results from the parsimonious models that increased substrate affinity decreases specificities. The PAA provides somewhat unnoticeable improvement over Ψ_4_ above the DUB switch threshold (*k_b_* > 0.003 s^-1^), whereas at the low DUB range Ψ_4_ becomes negative and the system prefers slow elongation to avoid PAA-induced strong E3-substrate binding at advanced conjugation states (Fig. 6e). The PAA further reduces substrate specificity when the minimal signaling chain length increases beyond *m* = 4 (see Fig. S9 for the *m* = 8 case). The specificity switches become more pronounced with PAA (Fig. 6e and Fig. S9e), because the PAA slows the stage-dependent decline in processivity and the system approaches closer to the unbounded elongation that has a sharp switch.

It is likely that the processive gain in substrate affinity to APC is a physicochemical phenomenon and could also be observed in nonsubstrates, in which case increased substrate-E3 affinity due to accumulation of ubiquitins is non-discriminative but speeds up ubiquitin transfers once a protein is initially conjugated. The elongation slowdown however renders the processivity gain less pronounced. It is also unclear whether the PAA is an APC specific or a rather universal phenomenon of substrate-ligase binding. Analysis of elongation kinetics of SCF substrates from single-contact bulk assays did not show stage-dependence of *k*_off_ [13]. Alternatively, the increased substrate degradation as observed in Lu *et al*. [3] could be attributed to increasing difference in ubiquitin transfer rates as ubiquitins accumulate because a substrate may expose more lysine sites to the donor ubiquitin. Indeed, human APC substrates on average have about 30% more lysine residues within a distance of 160 aa to the cognate D-box than nonsubstrates (9 vs. 7) [3], which may accelerate the substrate ubiquitin transfer. This hypothesis can be experimentally tested by the single E3-substrate contact assay that measures processivity or by direct measurement of individual ubiquitin transfer rates.

## Discussion

### Signaling efficiency-specificity tradeoff

Specificity and efficiency (or accuracy and speed) are two conflicting cellular goals. Improvement in one is often at the expense of the other. Their tradeoff is a fundamental and universal constraint on molecular recognition systems [24–26]. It remains elusive that what and how the cellular and molecular constraints architect the ubiquitination circuits to maintain a functional specificity-efficiency compromise. As suggested for the ribosome decoding, balancing efficiency and specificity may be resulted from tuning codon-anticodon binding affinity and the GTP activation rate to maximize a fitness function to favour recognition of cognate tRNAs over non-cognate or near-cognate ones [27]. The polyubiquitination is complicated by reversible substrate-E3 contacts, diverse modes of higher-order ubiquitin conjugations and DUB actions, and the stage-dependent kinetics. More fundamentally, proper efficiency versus specificity tradeoff may be constrained by limited cellular resources such as energy dissipation and by functional requirements such as signaling time.

Slow chain initiation was observed in ligase systems having either a single E2 (Cdc34 in SCF) or specialized E2s (Ube2C and Ube2S in APC) for chain initiation and elongation. Modulation of chain initiation rate was proposed to sensitively influence the signal strength [13] and control substrate degradation timing [28]. Slow chain initiation provides a low starting probability for substrate modification and a strong discrimination against nonsubstrates, approaching the Hopfield-Ninio limit of substrate binding free energy difference in the single-step Michaelis-Menten like mechanism. The chain initiation rate (*k*_*u*,0_ = 0.03 s^-1^) of the SCF substrate *β*-catenin is an order of magnitude slower than the dissociation rate constant (*k*_off_ = 0.3 s^-1^), corresponding to a probability of 0.1 to passage to the first conjugated state. A nonsubstrate with 10 times weaker E3 affinity (e.g., *k*_off_ = 3 s^-1^) has a conjugation probability 0.01. A slow chain initiation ensures that nonsubstrates of weak E3-binding are primarily rejected without ubiquitin conjugation, avoiding energy and material costs. Although any rate-limiting pre-signal elongation step can be as well exploited for substrate discrimination and can control signaling time, the slow initiation has a minimal energy cost. This suggests that energy dissipation could be a critical constraint that shapes the ubiquitination kinetics. Similar energy-efficient mechanics that engages slow initial processes to discriminate nonsubstrates discrimination was found in other molecular recognition systems such as tRNA recognition by the ribosome [11], where the GTPase activation for near cognate tRNA before GTP hydrolysis is markedly slow. The slow chain initiation is likely determined by structural difference between dedicated initiation and elongation E2s or by the relative inefficiency in searching for a substrate lysine site for systems using a single E2. In the system that uses a specialized initiation E2, slow initiation may be also achieved by a low E2 concentration.

Upon initiation, swift short-chain elongation allows efficient signaling and meanwhile provides additional proofreading to filter out nonsubstrates that bind to the ligase with affinity comparable to the substrates. The first ubiquitin conjugation constitutes a dominating step of substrate discrimination, while elongation events provide more opportunities to visit discriminative processes to enhance substrate selectivity at a reasonable cost of signaling efficiency (Fig. 4f and g). A 4-step ubiquitination of SCF substrate cyclin E provides a discrimination nearly 2 times the 1-step ubiquitination measured in specificities Φ_1_ = 0.45 vs. Φ_4_ = 0.79 and Ψ_1_ = 0.89 vs. Ψ_4_ = 1.61 (See Table S3). The self-avoiding walk model considers the elongation slowdown as a geometric limit as a ubiquitin chain spatially fluctuates to search for the constant reaction volume near the ubiquitin donor, rationalizing the scaffolding function of E3 ligase that structurally anchors the substrate and the E2 enzyme via their specific binding motifs. This contrasts other elongation processes such as protein translation or DNA replication, where ribosome or polymerase add monomeric base units with a length-independent rate. Since the elongation dissipates energy (at the E1-charging step) and requires ubiquitin recycling, the elongation slowdown prevents excessive polyubiquitination and offers an economical means to curb unnecessary energy and material expenditure after a substrate passes the minimal signaling state.

Substrate selection by the difference in ubiquitin transfer rates offers more rigorous proofreading than the substrate-E3 binding energy difference and may be a general mechanism in protein quality control by the ubiquitin system. Specifically, misfolded or damaged proteins may expose more ubiquitinable residues to facilitate chain initiation and elongation. By contrast, a folded protein may expose surfaces less accessible to ubiquitin conjugation and can only accumulate short ubiquitin chains subject to timely removal by DUBs. Similar kinetic disparities in substrate modifying steps (GTPase activation and tRNA accommodation into the A-site) were found between cognate and noncognate tRNAs and exploited to enhance fidelity of codon-anticodon match [11]. The discriminative role by the dissipative intermediate steps in molecular recoginition has been less discussed in the literature. In polyubiquitination, ubiquitin transfer events are difficult to directly observe and their rates can only be coarsely inferred from measurable quantities such as the ubiquitination processivity [13]. Single-molecule fluorescence techniques are promising and have the potential to resolve individual ubiquitin transfer rates [3].

### Minimal signaling chain length and linkage diversity

The minimal signaling chain length is another parameter that can be tuned to trade-off efficiency and specificity, as multiple steps in chain elongation reduce degradation signal but provide more proofreading. Given the elongation slowdown, the minimal signaling chain length is physically limited by the progressive loss of processivity (Fig. 4e, Fig. 6a and Fig. S9a) and therefore increasing minimal chain length accelerates the shift of the signal-specificity tradeoff towards high specificity and low signal (see Fig. S7 for simulations of SCF substrates cyclin E and *β*-catenin). An evolutionarily determined minimal chain length must seek a balance of efficiency and specificity at a reasonable energy, material and time costs. An adequate chain length (or ubiquitin count) should constitute motifs cognate to the proteasome. Large ubiquitin conjugates however require increased ubiquitin synthesis and burdens deubiquitination for ubiquitin recycle.

The Lys48-linked tetrameric conjugate has been widely-appreciated as the minimal ubiquitin complex to be recognized by the proteasome [29]. Various degradable chain length and linkages were also reported [30]. Instead of attacking a single lysine on its substrate to conjugate a ubiquitin chain, APC modifies a substrate by multiple short chains for more efficient recognition for proteasomal degradation [3, 31, 32]. Besides the often-studied homogeneous K11 or K48 linkage, heterogeneous chain linkages or branched networks have been found recognizable to the 26S proteasome [33]. Cellular functions implicated by such linkage complexity are yet to be fully characterized. Stepwise kinetics of ubiquitin conjugation for different linkages are even more difficult to resolve. Nonetheless, the model of sequential ubiquintin transfer is not restricted to study assembly of a single elongated chain and can be amended to model arbitrary linkages if suitable kinetics are identified. Linkage complication in higher-order ubiquitin complexes at least affects modeling of step-dependent ubiquitin transfer rates. Although the sequential ubiquitin transfer model does not rely the assumption of single ubiquitin chain, the self-avoiding walk model for the elongation slow down requires a polymer-like ubiquitin conjugate. Forming branched ubiquitin aggregates or multiple short chains originated from different substrate lysine residues implies that a donor ubiquitin could be conjugated to one of several residues on the acceptor ubiquitin complex. Such dynamic modulation of the transfer multiplicity as elongation progresses may cause a departure from the self-avoiding walk model that predicts the elongation rate as a monotonically decreasing function of the chain length. In any case, we expect that the elongation eventually slows down with the ubiquitin conjugate size as lysine residues on the conjugate grow in distance to the donor ubiquitin on E2 or sterically blocked.

On the other hand, the chain length and linkage dependence of DUB activities further complicates the modeling and interpretation. DUBs have diverse substrates and ubiquitin chain linkage specificity, which affects the structure of ubiquitination circuits and influences the signal and specificity tradeoff. Like the stage dependence of ubiquitin transfer and E3-substrate afinity, the deubiquitination rate may depend on structural accessibility of a substrate-ubiquitin or ubiquitin-ubiquitin isopeptide bond. Short Lys48 (mono- or diubiquitin) chains, for example, were found susceptible to DUB editing, while longer chains were more resistant to cleavage [34], suggesting that DUB actions beyond minimal degradable chain length were suppressed to increase signaling efficiency. A DUB may also cleave a bond within a ubiquitin chain, interpolating the two limiting cases of EBD and SQD.

### Sequential vs. *en bloc* ubiquitin transfer

Although sequential ubiquitin transfer as in SCF and APC are common [9, 13, 35], some E3-E2 systems transfer ubiquitins *en bloc* [36–38], in which a pre-assembled ubiquitin chain on E2 is transferred to a substrate in a single step. More realistically, the number of ubiquitins transferred per step is stochastic and depends on the chain length distribution on E2s. *en bloc* mechanism takes fewer intermediate steps for a substrate to be ubiquitinated into a signaling state and consequently lowers the step-dependent upper limit of specificities. Actual substrate specificity may not be substantially compromised as the discrimination is mainly implemented at the slow chain initiation and a ubiquitination system usually operates far below the theoretical limit achievable only at diminishing ubiquitin transfer rate. What is nonintuitive is that *en bloc* transfer does not necessarily increase processivity and ubiquitination efficiency as suggested [37]. Certain physiological conditions may even generate less degradation signal than the sequential transfer (Supplementary Materials). The processivity gain in bulk transfer is likely offset by a decrease in substrate conjugation rates caused by the longer waiting time in ubiquitin chain pre-assembly and the reduced availability of E2s conjugated with short chains, maintaining a relatively constant processivity.

### Error frequency in polyubiquitination

Although error rates in the ubiquitin-proteasome protein degradation remain unquantified to any E3 system, protein degradation is intuitively more tolerant to nonsubstrate degradation when compared to other high fidelity molecular recognition systems. Accumulated errors in amino acid or nucleotide cooperation during protein synthesis or DNA replication may cause critical cellular defects. By contrast, fortuitous degradation of a small fraction of a nonsubstrate population may not significantly alter its homeostasis and is likely balanced by a basal translation. Consider a decoy substrate binds to SCF 10 times weaker than *β*-catenin. It generates a degradation signal *S*_4_ about an unremarkable 1/60 of that of *β*-catenin (see Table S3 for the SQD model (*η* = 1)). By contrast, discrimination by difference in ubiquitin transfer rates provides a much more rigorous safeguard. A nonsubstrate with ubiquitin transfer rates 1/10 of *β*-catenin has the degradation signal is about 1/500 such that one nonsubstrate molecule degrades in parallel with 500 ligase-competing substrate molecules. This misubiquitination likely has little effect on biochemical pathways related to the nonsubstrate. Weak discrimination over substrate-binding energy potentially explains low specific ligase recognition motifs widely found in both substrates and nonsubstrates.

Accuracy of the ubiquitin-proteasome system (∼10^−2^ per molecule) is markedly lower than high-fidelity DNA replication (10^−10^-10^−8^ per nucleotide) and protein translation (10^−4^-10^−3^ per amino acid). This modest accuracy is not because of a weak proofreading in ubiquitination. The specificities Φ_4_ and Ψ_4_ are at least comparable to proofreading levels seen in tRNA decoding at the ribosome. The fidelity in tRNA recognition is first ensured by a large 1000-fold binding free energy difference between cognate and near cognate tRNAs [24]. Binding free energy and ubiquitin transfer differences between a true substrate and its lookalikes are often less than 10 fold. The error rate of 10^−4^ per aa-tRNA incorporation implies that 1 in 25 proteins of average size (400 aa) may contain one misincorporated amino acid, which is comparable to the above error rate of nonsubstrate degradation by polyubiquitination on per protein basis. *in vitro* processivity measurement showed that only 6.4% *β*-catenin were polyubiquitinated into signaling states after a single-contact with SCF [13]. The steady-state level could be even lower when DUB activity is considered (below 1% when *k_b_* = 1 s^-1^, Fig. 4g). Low degradation signal further suppresses more than haphazard nonsubstrate degradation.

### Degradation rate and temporal ordering

The degradation rate or half life of a substrate is determined by: (i) the steady-state ubiquitination level (or equivalently, the processivity), when protein destruction at the 26S proteasome is rate-limiting; or (ii) the ubiquitination kinetics, when the proteasomal degradation rate is comparable to substrate ubiquitination or a slow ubiquitination becomes a bottleneck to substrate degradation. Different E3 enzymes coordinate ubiquitination in different kinetics. APC substrates, for example, have slow dissociation (*k*_off_ = 0.01 s^-1^) [39, 40] and are ubiquitinated in minutes (*k_u_* ∼ 0.1 s^-1^) [3, 9, 41]. SCF substrates, by contrast, dissociate an order of magnitude faster (*k*_off_ = 0.3 – 0.4 s^-1^) and are ubiquitinated in seconds (*k_u_* ∼ 5 s^-1^) [13] due to rapid E2 Cdc34 binding and dissociation [42]. Despite large kinetic disparities across ligase systems, the processivity appears to be relatively invariant (Fig. 4e and Fig. 6a), presumably to balance signal and specificity at an optimal tradeoff. This requires that E3-substrate binding, ubiquitination, and deubiquitination rates are in a same time scale (as shown in Eq. 1, *ρ_n_* stays invariant when all kinetics vary by a same factor). The steady-state degradation signal *S_m_* is related to the processivity (see Eq. S10 for the parsimonious SQD model), while the rate of signaling depends on kinetics as a fast circuit produces a quick signal [43].

During the metaphase to anaphase transition in the cell cycle, APC ubiquitinates cyclins and cyclin-dependent kinase inhibitors for proteasomal degradation in a meticulous order. How does APC favourably recognize a certain subset of substrates for polyubiquitination over others in time? This question is especially relevant when substrates are controlled by the same activator protein. The question may be addressed in the same principle as in how a ligase distinguishes substrates from nonsubstrates. It has been proposed that substrate processivity may determine the timing of degradation [9, 44], as high processive substrates are degraded earlier than low processive (distributive) ones. As the cellular concentration of proteasomes is limited (∼8x 10^5^ /cell [45]), we expect that proteasomes function near the saturation level while a large number of candidate substrates compete for a small population of proteasomes. Under this circumstance, substrates with higher steady-state degradation signal *S_m_* occupy more proteasomes and thus degrade faster. Unlike the graded responses to varied DUB activities in the EBD model, small changes in DUB activity in the SQD model can shift ubiquitination from a restricted elongation regime to an extensive elongation regime, or vice versus, which acts as an effective switch to control substrate specificities (Fig. 3 and Fig. 4). Below a threshold of the DUB activity, substrates and nonsubstrates are non-discriminatively ubiquitinated for degradation. At a mild cost of ubiquitination efficiency, increasing DUB acitivity above the threshold turns on the proofreading mechanism that rejects nonsubstrates for polyubiquitination. DUB activity substantially higher than the threshold however also inhibits substrate ubiquitination. This suggests that the active DUB concentration should be regulated (through phosphorylation or degradation) within a narrow functional range. The DUB threshold for a specific substrate is modulated by its kinetic parameters (or processivity. See Eq. S18 for the separation of bounded and unbounded elongation regimes in the parsimonious SQD model). The importance of DUBs in APC-mediated substrate degradation ordering has been demonstrated [9], where DUB stabilizes low-processive substrates while allows high-processive substrate to be polyubiquitinated for proteasomal degradation. Our results show that the ultrasensitive switch-like responses to the DUB activity could be potentially utilized to establish a temporal degradation degradation order of substrates with small processivity differences, by progressively deactivating DUBs, which allows ubiquitination of substrates in an order according to their descending processivity 7. Or conversely, the ubiquitin transfer rates (thus the processivity) can be progressively tuned up (by increasing ubiquitin-conjugated E2s) to match the deubiquitination to establish the polyubiquitination order in substrates. Free active ligase concentration can be another factor that regulates the switch, which controls the probability ratio of rebinding to deubiquitination (or the backward processivity). Degradation of high processive substrates reduces the competition for ligases and increases its availability to existing substrates. Even though increasing free ligase concentration does not alter the processivity, it does lower the probability of backtracking and thus allows low processive substrates to be polyubiquitinated. Sequential ubiquitination and deubiquitination circuit in structure resembles multisite phosphorylation in signal transduction where switch-like responses were common [46, 47], suggesting that higher-order (multisite or elongation) post-translational modification could be a general strategy in regulating on-and-off cellular decisions.

**FIG. 7.**
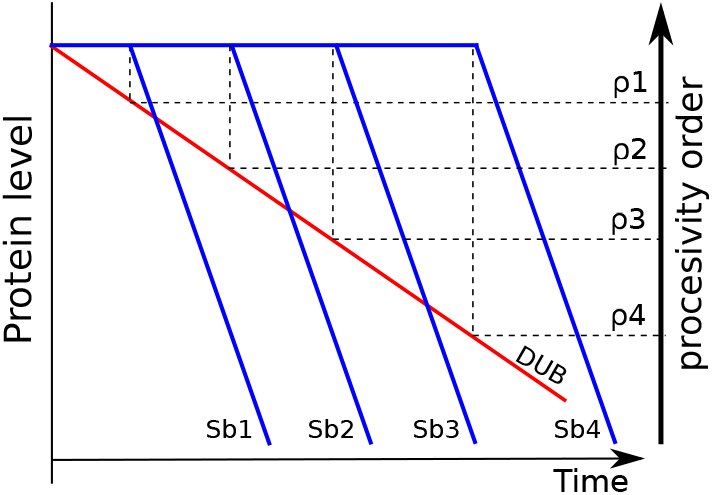
Temporal order of degradation. Substrates Sb1, Sb2, Sb3, and Sb4 have different processivities (*ρ*_1_ > *ρ*_2_ > *ρ*_3_ > *ρ*_4_). Slowly decreasing DUB activity (red), or equivalently the backward processivity, establishes a temporal order in the substrates (blue) as higher processive substrates degraded earlier. The time axis is shown in a logarithmic scale.

## ACKNOWLEDGEMENT

This study was supported by NSFC grant 308070477 and SIBS-Sanofi Innovation Grant to JY. We thank Xin Meng, Hong Zhang, Wenhong Hou, Weiren Cui, and John Pearson for helpful discussions.

## Supplementary Materials – Mathematical Derivations and Additional Results

Here, we first use the classic Michaelis-Menten reaction as the example to introduce the concept and definition of specificity. We then detail the mathematical derivations of the parsimonious EBD and SQD model and numerical computation methods used to simulate models with nonuniform parameter values. Extra results, including supplementary tables and figures (referred in the main text), are also presented.

### Rate and specificity of Michaelis-Menten reaction

In this section, we introduce the definition of specificities and their calculation using the following Michaelis-Menten model:

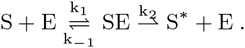

Assuming the complex SE is at a (pseudo)steady state and the free enzyme concentration is a constant, the probability *p* for a substrate molecule being in the intermediate complex is determined by *k*_1_ [E](1 – *p*) = (*k*_-1_ + *k*_2_)*p*, and therefore

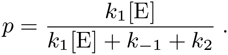

The enzyme *specificity* is conventionally defined by the ratio between conversion rates of substrate *S* and nonsubstrate (or near substrate) *S*’ of equal concentration:

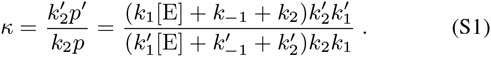

The lower *κ* is, the higher the specificity. When the binding rate constants of the two substrates are comparable and conversions are slow, i.e., 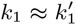 and 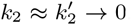. At the cost of the production rate, *κ* approaches the minimum:

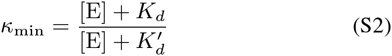

At low enzyme concentration 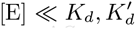, it recovers the Hop-field limit 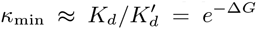, where Δ*G* = *G*’ – *G* is the binding energy difference (in units of *k_B_T*) between *S*’ and *S*. We note that, in Hopfield’s treatment [4], the free enzyme concentration [E] does not appear as in the above because both free substrate and free enzyme concentrations were assumed equal for the two substrates. Instead, we derive Eq. S1 by comparing conversion rates of two single molecules of the substrate and the nonsubstrate under a constant pool of enzyme molecules. (equivalent to the assumption that total concentrations of the two substrates are equal and they compete for enzyme binding sites).

A much less discussed branch of substrate discrimination is the difference in kinetic transition barriers (*k*_2_ and 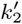. If the enzyme binding free energy difference between the substrate and nonsubstrate is small 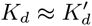, Δ*G* ≈ 0, we have

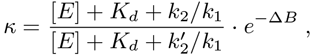

where Δ*B* = *B*’ – *B* is the difference of transition energy barriers. For large *K_d_* (Again, at the cost of production rate), the specificity approaches a limit:

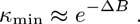

**FIG. S1.**
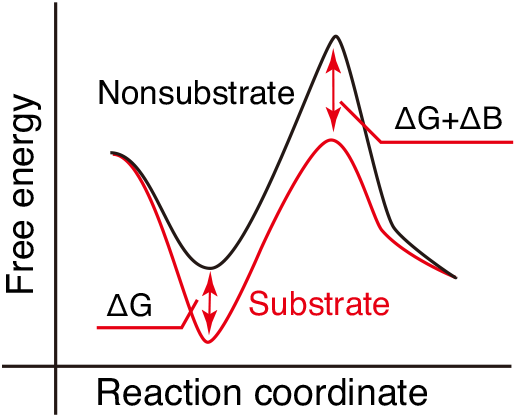
Free energy landscape of Michaelis-Menten reaction. A substrate (red) and a nonsubstrate (black) are distinguished by the enzyme binding free energy difference Δ*G* and the transition energy barrier difference Δ*B*.

In combination, the Michaelis-Menten reaction approaches the maximum specificity at the limit of weak binding and low conversion:

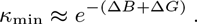

The free energy landscapes shown in Fig. S1 illustrate the binding energy and transition barrier differences between a substrate and a nonsubstrate.

As an alternative to the production ratio in Eq. S1, we define the enzyme specificity: the relative sensitivity of the catalytic rate to a local change of a kinetic parameter. This definition has the advantage of working with one set of parameters for a given model without explicitly comparing production rate of a substrate to that of a nonsubstrate. The normalized rate *υ* ≡ *k*_2*p*_/(*k*_1_[E]) can be written as a function of dissociation rate constant and conversion rate constant:

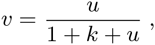

where *k* ≡ *k*_-1_/(*k*_1_[E]) and *u* ≡ *k*_2_/(*k*_1_[E]). The conversion rate *υ* is limited by the substrate-enzyme binding rate *k*_1_ [E] (*υ* < 1). The total logarithmic difference of *υ* with regard to *k* and *u* is:

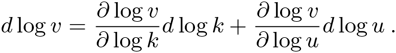

The specificity over enzyme-substrate binding free energy under a constant conversion rate *u* is now defined and calculated as:

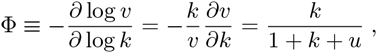

which measures orders of magnitude change in v per order of magnitude change in *k* (local to the value of *k*). We note that Φ can be related to the conventional rate ratio between substrates *S*_1_ and *S*_2_ by *υ*_2_/*υ*_1_ ≈ (*k*_2_/*k*_1_)^-Φ^ when the difference between *k*_2_ and *k*_1_ is small. The larger Φ is, the higher the specificity. For small *u*, Φ ≈ *k*/(1 + *k*), ranging from 0 to 1 as *k* varies. Within the tight binding regime (*k* ≪ 1), the enzyme specificity decreases as Φ approaches 0. In contrast, a weaker binding promotes specificity (i.e., *k* → ∞, Φ → 1) but reduces the conversion rate *υ*, showing a tradeoff between *υ* and Φ.

The enzyme specificity over the conversion rate *u* under a constant binding affinity *k* is defined as:

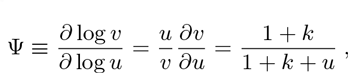

which is limited by 1 at large k and reaches the lower limit 1/ (1 + *u*) when *k* → 0. A substrate with a smaller *u* has greater specificity as Ψ is a monotonically decreasing function of *u*. More importantly, the specificity over the conversion rate is consistently larger than that over substrate binding energy, Ψ – Φ = 1/(1 + *k* + *u*). The relationship Ψ > Φ holds for more general kinetic proofreading schemes as we will show in the ubiquitination models.

### Ubiquitination models

Here, we present mathematical details of the kinetic ubiquitination models and derive analytical formulas to essential steady-state quantities for the parsimonious EBD model (Fig. 1b) and SQD model (Fig. 1c) that are parameterized by stage-independent (uniform) dissociation, ubiquitination and deubiquitination rates.

*en bloc* **deubiquitination (EBD).** We normalize the rate constants by the substrate-E3 binding rate, *k*_+_ = *k*_on_[E3], such that *u* = *k_u_*/*k*_+_,*k* = *k*_off_/*k*_+_, *b* = *k_b_*/*k*_+_, and *τ* = *k*_+_*t*. Consider a single substrate molecule interacts with a fixed population of E3 ligase molecules. The master equation for the system is written as:

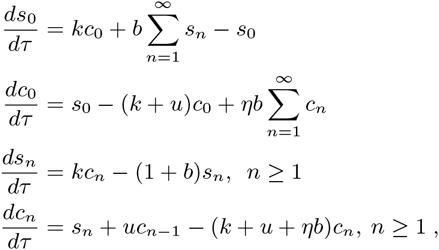

where *c_n_* or *s_n_* are probabilities of the substrate in an E3-associated or off-E3 state with *n* ubiquitins conjugated. Without explicitly considering substrate degradation, the total probability is conserved, 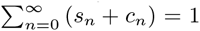. To aid interpretation, we define three probabilities:

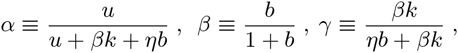

where *α* is the probability for an E3-bound and modified substrate *c_n_* to be incrementally modified to state *c*_*n*+1_ without backtracking. *β* is the probability for a dissociated and modified substrate *s_n_* to be deubiquitinated to the basal state *s*_0_, and *γ* is the probability that *c_n_* is deubiquitinated via the substrate dissociation pathway. The steady-state fractions *c_n_* and *s_n_* are geometric series:

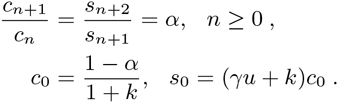

Both *c_n_* and *s_n_* are convergent since *α* < 1.

### Sequential deubiquitination (SQD)

The system of sequential deubiquitination is described by the following master equation:

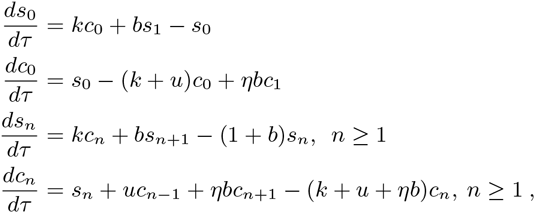

again, under the probability conservation: 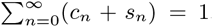. Alternative steady-state equations (not independent from the above master equation) can be derived to replace one of last two equations in the above, by accounting for in and out fluxes of a sub network (instead of balancing fluxes through a state).

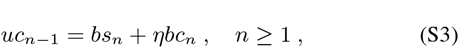

by the fact that the ubiquitin transfer rate from *c*_*n*-1_ to *c_n_* is balanced by deubiquitination rate from *s_n_* to *s*_*n*-1_. We give explicit steady-state solutions to the case of *η* = 0. The case of *η* = 1 relies on numerical simulations. At the steady state, these equations lead to a geometric ratio:

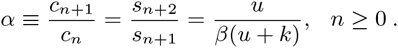

The system undergoes bounded elongation only when *α* < 1. Given *bs*_1_ = *uc*_0_ from Eq. S3 when *η* = 0, we have:

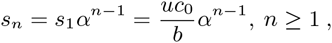

where

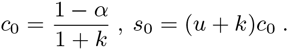

When *α* > 1, the system undergoes an unbounded elongation.

In the following sections, we derive explicit results to chain length statistics, processivity, degradation signal, and specificities. Results in analytical formulas are also summarized in Table S1.

### Chain length distribution, mean chain length and variance

Chain length statistics can be obtained from the stationary chain length distribution *w_n_* For the EBD model,

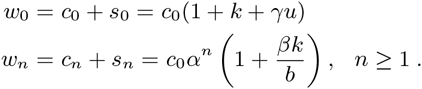

The mean chain length serves as a state-independent measure of the overall ubiquitination level,

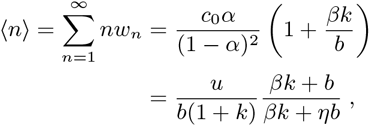

which is proportional to the ubiquitin transfer rate and limited by E3-substrate affinity and the deubiquitination rate. Without E3-bound DUB editing (*η* = 0), 〈*n*〉 is limited by the substrate dissociation rate when *b* is large. With E3-bound DUB editing (*η* = 1), 〈*n*〉 is inversely proportional to *b*. The mean chain length of dissociated substrate *s_n_* is:

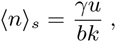

The variance of chain length is

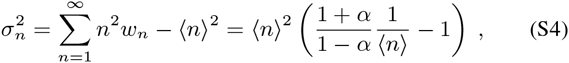

which applies to both 〈*n*〉 and 〈*n*〉_*s*_. The fluctuation of chain length around the mean is then

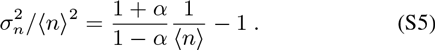

One can verify that the fluctuation 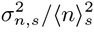 is substantial (> 1)and can be tuned arbitrarily large, especially when *α* is small.

**FIG. S2.**
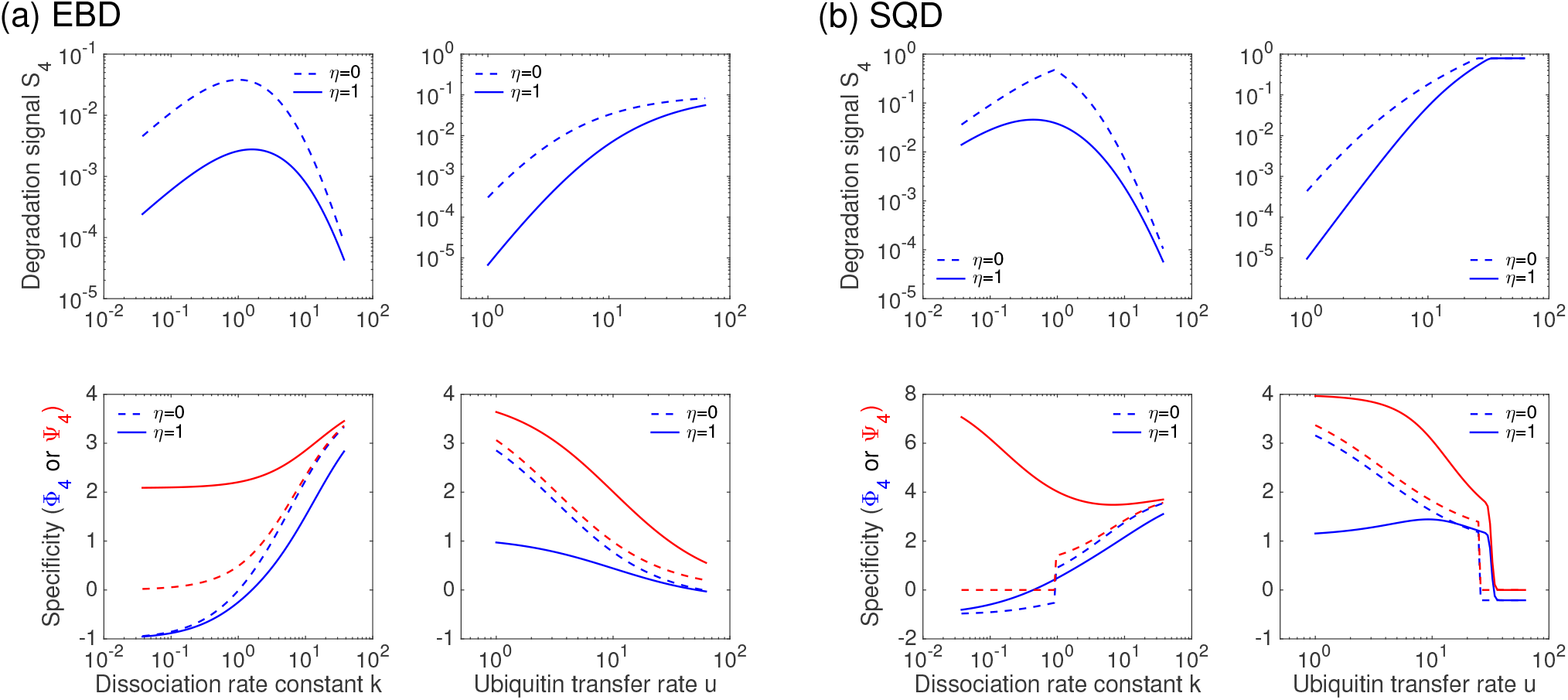
Steady-state degradation signal and specificities in the EBD and SQD models (complementary to Fig. 2). *S*_4_, Φ_4_, and Ψ_4_ vs. *k* and *u*. Parameter values are described in Fig. 2. Quantitatively, the specificities Φ_4_ and Ψ_4_ measure the slopes of *S*_4_ vs. *k* and *u* curves, respectively. Consequently Φ_4_ and Ψ_4_ could over- or under-estimate responses of *S*_4_ to large changes in *k* and *u* (see Table S3 for comparison between small and large parameter changes.). Processivity changes in a narrow range could quickly switch the specificities up and down in the SQD model.

For the SQD model of bounded elongation, when *η* = 0 the mean chain length for all substrates and the dissociated substrates are:

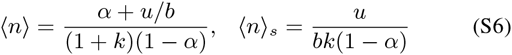

The variance formula of Eq. S4 applies to the SQD model as well.

### Processivity *ρ_n_*

The mean chain length conjugated to a substrate after a single E3 contact defines the ubiquitination *processivity*, a measurable quantity in a substrate-E3 single-encounter assay [13, 48]. The processivity *ρ_n_* depends on the starting state on the E3, *c_n_*. Below, we derive a general formula to *ρ_n_* for the EBD model. Consider transition probabilities of an E3-bound substrate taking alternative reaction paths (ubiquitination, dissociation, or deubiquitination) as:

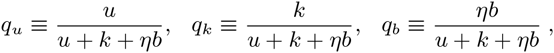

with *q_u_* + *q_k_* + *q_b_* = 1. To calculate *ρ_n_*, we first determine the probability *p*_0*l*_ that a substrate dissociates from the E3 ligase at state *c_l_* when it starts its stochastic transitions at *c*_0_. *p*_0*l*_ accounts for probabilities via two pathways that lead to a dissociation event from *c_l_*: (i) a walk directly to *c_l_* for the first time by ubiquitination and then dissociates without looping back to *c*_0_, and (ii) random walks to any state *c_i_*, *i* > 0 and then backtrack to *c*_0_ (due to deubiquitination), where the probability *p*_0*l*_ applies again:

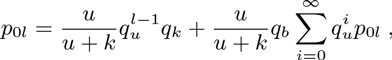

from which *p*_0*l*_ can be derived:

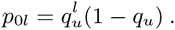

The above formula also applies when *l* = 0. We then determine the more general probability distribution, *p_nl_*, for a substrate to dissociate from state *c_l_* given it starts at state *c_n_*. *p_nl_* also accounts for probabilities of two pathways from *c_n_* to a dissociation event at state *c_l_*: (i) a walk from *c_n_* to *c_l_* when *l* ≥ *n* for the first time without deubiquitination and then dissociates from the E3, and (ii) a random walk from *c_n_* back to *c*_0_ via deubiquitination en route any downstream state *c_i_*, *i* ≥ *n*, where *p*_0*l*_ applies again:

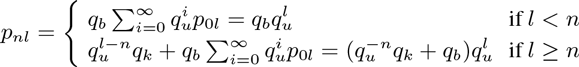

The processivity reaches a rather simple equation:

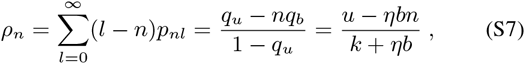

which is regulated by all kinetic key parameters. Without E3-bound DUB editing, *ρ_n_* = *u*/*k*, independent of *n*. With E3-bound DUB editing, *ρ_n_* linearly decreases with *n*, indicating that rebinding of a sufficiently conjugated substrate to the E3 ligase does not contribute to further elongation. *ρ_n_* becomes negative (backward processivity) when *n* > *u*/*ηb*, where E3-bound deubiquitination outcompetes ubiquitination. We can also calculate the fluctuation of processivity:

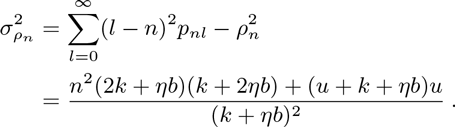

For the SQD model when *η* = 0, the processivity is state-independent, *ρ* = *u*/*k*, like in the EBD model. When *η* = 1, *ρ_n_* can be calculated by numerical simulations, by integrating ODEs of the model with *s_n_*, *n* = 0, 1,…, as absorbing states under the initiation condition *c_n_* = 1, *n* = 0,1,…, until probabilities are all absorbed to obtain the distribution *p_nl_*. This method is generally applicable to models of nonuniform rates.

**TABLE S1.** Formulas to key quantities in parsimonious EBD and SQD (*η*=0) models

| Model | MCL $\langle n \rangle_s$ | $\sigma_n^2 / \langle n \rangle^2$ | Processivity $\rho_n$ | Signal $S_m$ | Specificity $\Phi_m$ | Specificity $\Psi_m$ |
| --- | --- | --- | --- | --- | --- | --- |
| EBD | $\frac{\gamma u}{bk}$ | $\frac{1+\alpha}{1-\alpha} \frac{1}{\langle n \rangle} - 1$ | $\frac{u - \eta bn}{k + \eta b}$ | $\frac{\alpha_e^m \beta k}{(1+k)b}$ | $m(1-\alpha_e)\gamma - \frac{1}{1+k}$ | $m(1-\alpha_e)$ |
| SQD ($\eta=0$)* | $\frac{u}{bk(1-\alpha_s)}$ | $\frac{1+\alpha}{1-\alpha} \frac{1}{\langle n \rangle} - 1$ | $\frac{u}{k}$ | $\frac{u\alpha_s^{m-1}}{(1+k)b}$ | $\frac{(m-1)k}{u+k} + \frac{k}{1+k}$ | $\frac{(m-1)k}{u+k} + 1$ |

$\alpha_e = u/(u + \beta k + \eta b)$ for the EBD model or $\alpha_s = u/(\beta(u + k))$ for the SQD model.
$\beta = b/(1+b)$ and $\gamma = \beta k/(\beta k + \eta b)$ .
\* Quantities for the SQD model when $\eta = 1$ are obtained by simulations.

### Degradation signal *S_m_*

Dissociated substrates conjugated with ubiquitin chains of at least the minimal length *m* are recognizable to the 26S proteasome, which is defined as the degradation signal in our study. The degradation signal in the EBD model is calculated as:

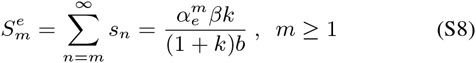

and the degradation signal in the SQD model (*η* = 0) is:

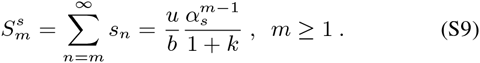

One can verify that in all cases 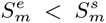. Eq. S9 applies only when the chain elongation is bounded. In this case, the degradation signal and the processivity (*ρ* =*u*/*k*) can be related by the following

**FIG. S3.**
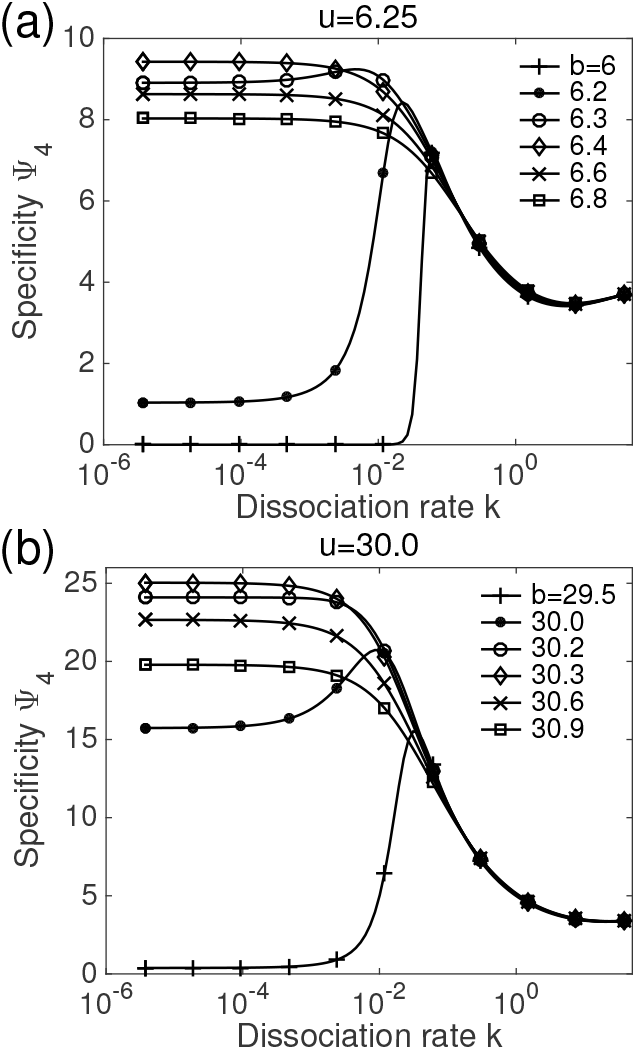
Ultra-high discrimination by Ψ_*m*_ in the SQD model of E3-bound DUB editing. (a) Specificity Ψ_4_ vs. *k* under varying *b* around *u* = 6.25. Other parameters are specified in Fig. 2 and (b) *u* = 30. Ψ_4_ reaches a maximum plateau at small *k* when *b* is slightly larger than *u* (e.g., *b* = 6.4 when *u* = 6.25), allowing the biased random walk along the elongation path to returns to presignaling state, *c_n_* < *m*, before substrate dissociation.

relationship when *k* is larger than 1:

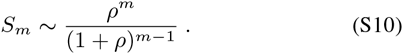

When the elongation is unbounded, all substrates molecules are eventually polyubiquitinated and the degradation signal is determined by the E3-substrate binding equilibrium, i.e., *S_m_* = *k*/(1 + *k*) that is independent of ubiquitin transfer and deubiquitination and increases with k as the processivity decreases.

### Specificities Ψ*_m_* and Ψ*_m_*

The logarithmic total derivative of the degradation signal is

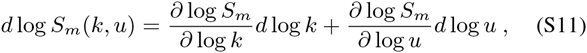

and the substrate specificity is defined as sensitivity of *S_m_* to binding affinity *k* or ubiquitin transfer rate *u*:

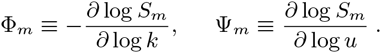

The minus sign ensures that a positive Φ*_m_* indicates discrimination against a weaker binding substrate.

For the EBD model, we have:

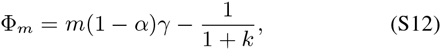

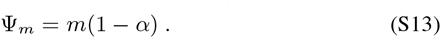

These equations provide fast insights to the specificities of the EBD model, which in general hold for systems with experimentally characterized kinetic rates. Φ*_m_* can be interpreted as the sum of probability for a substrate of state *c_n_*, *n* < *m* to be deubiquitinated to basal state so via the substrate dissociation pathway up to stage m (the first term), which is offset by the sequestration effect by the E3 (the second term). The degradation signal *S_m_* attains a maximum when Φ*_m_* =0 as *k* approaches a non-discriminative value:

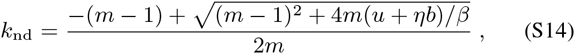

which increases with *u* and decreases with *b* when *η* = 0, and has a non-monotonic relationship with *b* when *η* = 1. The limiting behaviours are independent of model choices:

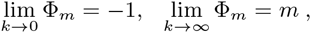

which encloses a two-regime proofreading as the E3-substrate affinity *k* varies: (1) the model proofreads against tight binding (–1 < Φ*_m_* < 0 when *k* < *k*_nd_) or (2) against weak binding (0 < Φ*_m_* < *m*

**FIG. S4.**
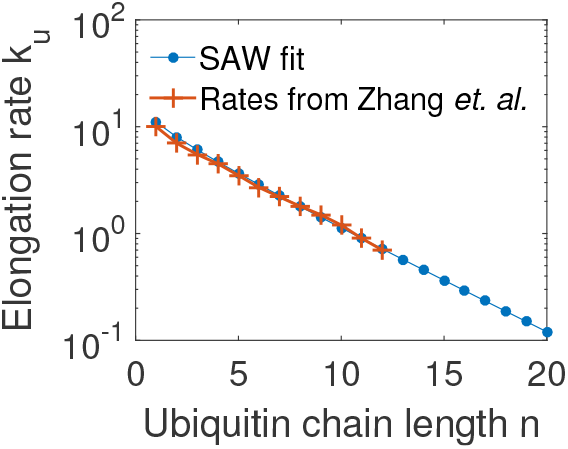
The self-avoiding walk model fit (*k*_*u*,*n*_ = 13.68 × 1.238^-*n*^ × ra^-0.157^ s^-1^, blue curve) to the ubiquitin transfer rates (red markers) used by the model in Ref. [10].

when *k* > *k_nd_*). The limiting behaviors when the ubiquitin transfer rate *u* changes are given as:

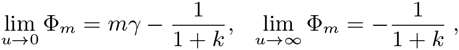

By comparison, specificity Ψ*_m_* accounts for the sum of probability for a substrate to be deubiquitinated to the basal state *c*_0_ or *s*_0_ via either the substrate dissociation or the direct E3-bound DUB pathway. Ψ*_m_* is a positive and monotonically decreasing (or increasing) function of ubiquitin transfer rate *u* (or dissociation rate *k*), with limiting cases:

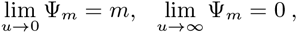

and

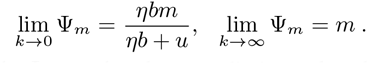

In other words, Ψ*_m_* reaches the upper limit *m* when the processivity *ρ* approaches zero and diminishes at large processivity *ρ*.

The specificities for the SQD model when *η* = 0 are given as:

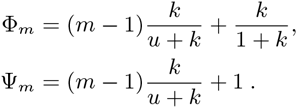

Fortuitously for this case, because terms containing *b* in *S_m_* are multiplicatively separable from other parameters, both specificities are independent of the deubiquitination. The difference between Ψ and Ψ*_m_* is a constant. For the case of *η* = 1, Ψ*_m_* and Ψ*_m_* are computed numerically.

The finite-difference form of Eq. S11 can be used to estimate energetic or kinetic difference between a substrate and a nonsubstrate. As an example, we show how to use Eq. S11 to predict the difference in ubiquitin transfer rates of SCF substrate CD4 and its mutant CD4-M. The estimation is based on known data and calculated specificities. The study by Zhang *et al*. [10] found that DUBs amplified the difference in polyubiquitinations of CD4 and CD4-M to 5-10 fold from a modest difference in their affinities to Vpu 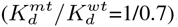. Using the SQD model with E3-bound DUB editing, we calculated Φ_4_ = 1.17 and Ψ_4_ = 1.59 at *k_b_* = 0.5 s^-1^(see the Main Text). We assume that the ubiquitin transfer rates of a substrate and a nonsubstrate are related by a scaling factor *w* (identical in each individual rates), i.e., 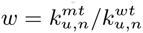. Using the formula

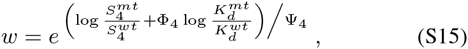

we can predict that the ubiquitin transfer rates of CD4-M are about 31–47% of the wild type CD4.

### Effects of deubiquitination

In all model variants, *S_m_* decreases as DUB activity increases with the limiting cases: (i) lim_*b*→0_ *S_m_* = *k*/(1 + *k*), where the substrate elongates indefinitely and equilibrates between the unbound and bound forms, and (ii) lim_*b*→∞_ *S_m_* = 0. In reality, the deubiquitination rate is limited by *k*_cat_, which can be approached when DUBs saturate the substrates.

*EBD model*. In the EBD model when *η* = 0, for small *b*,

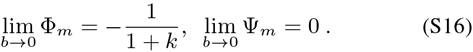

All substrates and nonsubstrates are indiscriminate polyubiquitinated, and the system favors weaker E3-binding substrates and is insensitive to differences in ubiquitin transfer. When *b* ≫ 1, the system effectively allows only one E3-substrate encounter before a deubiquitination event. The ubiquitin chain is instantly removed from the substrate by the DUB as soon as a substrate dissociates from the E3. In this regime, the system approaches the Hopfield-Ninio model.

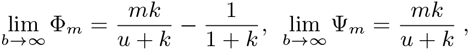

indicating that increasing processivity reduces specificity and shifts the model to proofread against strong binding.

When *η* = 1, for small *b*, the model converges to the *η* = 0 case in Eq. S16. As the DUB rate *b* increases, both specificities Φ*_m_* and Ψ*_m_* increase. But they diverge when the DUB activity *b* > *b*_opt_, with the limiting behaviours:

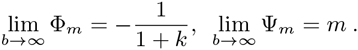

Φ*_m_* attains a maximum when *∂Φ_m_*/*∂b* = 0 at the optimal deubiquitination rate *b*_opt_ = *u*^1/2^, and

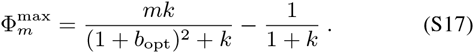

*SQD model—Bounded and unbounded elongation*. An important dif-ferencebetweenSQD andEBDmodels is thatthechainelongationof the former may not be bounded under some parametric conditions, whereas that of the latter is always bounded because of the single-step backtracking to the basal state to prevent a runaway elongation. When *η* = 0, switched by the substrate binding and unbinding with the E3 ligase, the SQD model has a symmetry in forward random walk along the ubiquitination pathway and backward random walk along the deubiquitination pathway (Fig. 1). The forward processivity is *ρ* = *u*/*k* and the backward processivity is *b*. The backward processivity of a bounded elongation must be larger than its forward processivity:

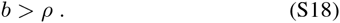

When *ρ* > *b*, the model has an unbounded elongation. The degradation signal can be obtained by the simple argument that steady-state *C_m_* and *S_m_* are at the binding equilibrium such that *S_m_* = *k*/(1 + *k*). The specificities can be therefore calculated:

**TABLE S2.** Signals ofthe CD4-Vpu-SCF system(Fig. 5)

| Substrate | $k_{\text{off}} (\text{s}^{-1})$ | $w$ | $S_4$ | $S_4/S_4^{\text{wt}}$ |
| --- | --- | --- | --- | --- |
| 1 (WT) | 0.4 | 1 | 0.1211 | 1 |
| 2 | 0.6 | 1 | 0.071 | 0.59 |
| 3 | 0.4 | 0.5 | 0.034 | 0.28 |
| 4 | 0.6 | 0.5 | 0.018 | 0.14 |
| $\Phi_4 = 1.17$ and $\Psi_4 = 1.59$ for wild-type substrate | | | | |
| $k_b = 0.5 \text{ s}^{-1}$, $k_{\text{on}} = 0.01 \text{ nM}^{-1} \text{ s}^{-1}$, and $[\text{E3}] = 8 \text{ nM}$. | | | | |

**FIG. S5.**
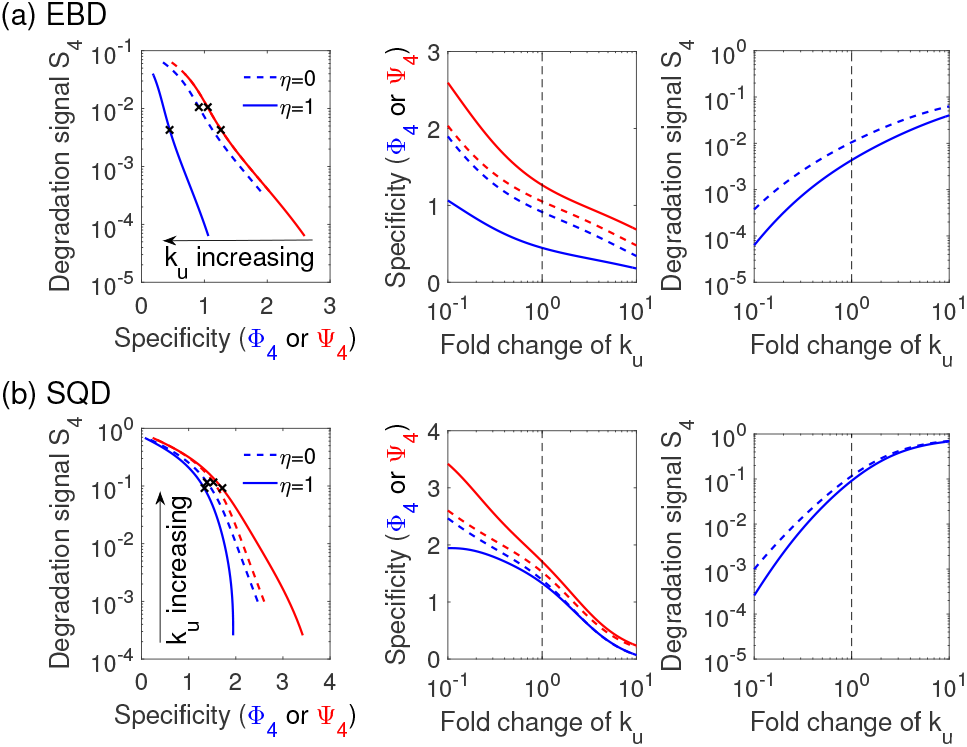
Effects of ubiquitin transfer rates on specificities and degradation signal. (a) EBD and (b) SQD model. Ubiquitin transfer rates *k_u,n_* at all conjugation stages vary by an identical proportion from one tenth to 10 fold of the nominal rates *k*_*u*,0_ = 0.05 s^-1^ and *k*_*u,n*_ = 13.68× 1.238^-*n*^ × *n*^-0.157^ s^-1^ that extrapolate rates from Ref. [10] (Fig. S4). *S*_4_, Φ_4_, and Ψ_4_ calculated from nominal *k*_*u,n*_ ‘s are marked (’x’ in the two left panels and vertical dashed lines in the middle and right panels). *k*_off_ = 0.4 s^-1^, *k_b_* = 0.5 s^-1^, *k*_on_ = 0.01 nM^-1^s^-1^, and [E3]=8 nM.

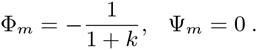

This sharp transition between bound and unbound elongation also exists for the case *η* = 1. Because of E3-bound DUB editing, the specificity switch has a DUB threshold lower than that in Eq. S18 for *η* = 0 (Fig. 3a).

### Chain initiation

Polyubiquitination by an E3 takes two steps, chain initiation and elongation, catalyzed by different E2s in many cases. The chain initiation is the step by which the E3-E2 pair conjugates the first ubiquitin molecule to the substrate and has been considered typically slower than the subsequent elongation. We show below the influence of chain initiation on degradation signal and substrate specificities. We assume that the initiation proceeds at rate *u*_0_ and elongation at a state-independent rate *u* (typically, *u*_0_ < *u*).

For the EBD model, we modify the differential equations that describe the dynamics of *c*_0_ and *c*_1_:

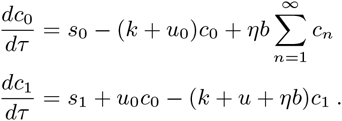

Define the ratio *λ* = *u*_0_/*u*. We derive:

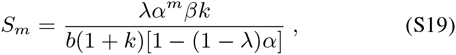

which is the degradation signal of the parsimonious (uniform elongation rate) model multiplied by a factor *λ*/(1 – (1 – *λ*)*α*) that increases with *λ*. When *λ* < 1, the chain initiation is slower than the subsequent elongation and the degradation signal is reduced, in comparison to that in the model of uniform ubiquitin transfer rates.

When *λ* = 1 (i.e., *u*_0_ = *u*), the model recovers the parsimonious model. When *λ* > 1, the processivity *ρ*_0_ increases over the parsimonious model and so does the degradation signal. We can calculate the specificities:

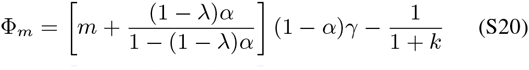

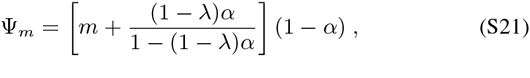

which again show the tradeoffs with *S_m_* as *λ* varies. We can alternatively write the equations of *S_m_* and specificities to assist interpretation. Let *α*′ = *u*_0_/(*u*_0_ + *βk* + *ηb*). One can verify:

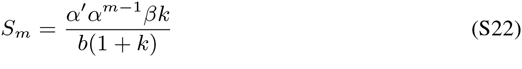

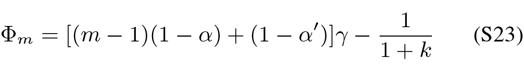

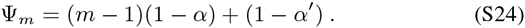

Compared to equations to the parsimonious models (Eqs. S8, S12, and S13), each of the above equations replaces one *α* component (either a multiplicative component in *S_m_* or an additive one in specificities) with *α*′.

For the SQD model (*η* = 0), we need two more steady-state equations:

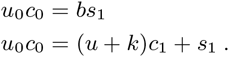

We can derive:

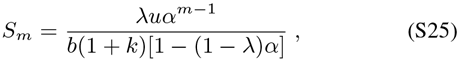

which relates to the parsimonious model by a same factor as that in the EBD model. The two specificities are calculated as:

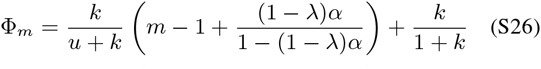

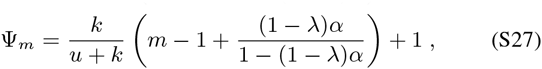

which are not independent of DUB activity. In general, any pre-signal rate-limiting ubiquitin transfer *u_n_*, *n* < *m* would have a dominant influence on E3 specificity, signal magnitude, and the timing of degradation. On the contrary, a faster chain initiation rate (i.e., *λ* > 1) increases degradation signal at the expense of specificities.

### Self-avoiding walk (SAW) model of elongation slowdown

The E3-mediated ubiquitin transfer can be viewed as a searching process in which the acceptor ubiquitin at the distal end of a ubiquitin chain fluctuates in space and stochastically locates near the donor ubiquitin on E2. This process can be approximated by the SAW model originally developed to model polymer growth. The SAW model enumerates all distinct non-self crossing configurations for a ubiquitin chain of a given length *n* on a lattice (see Fig. 4 for a 2D illustration) and relates the ubiquitin transfer rate *u_n_* to the probability for the acceptor ubiquitin to fluctuate near the donor. *u_n_* therefore decreases with the chain length as the total searching volume grows when *n* increases and the reactive volume remains relatively unchanged, in agreement with observations of the elongation slowdown [13].

Given the above consideration, we apply established results from the SAW model to extrapolate experimental data. The number of configurations for a chain of length *n* can be approximated as *N*_saw_ ∼ *Aμ^n^n^a^* [49]. The reactive volume that encloses the donor ubiquitin is limited by a fixed size and therefore corresponds to a fraction of the total number of chain configurations. Assume that each chain configuration has an equal probability and this reactive volume is much smaller than the whole SAW volume. The probability that a ubiquitin chain is located inside the reactive volume is (roughly) inversely proportional to *N*_saw_. We use the following equation to generate ubiquitination rates *u_n_*:

**FIG. S6.**
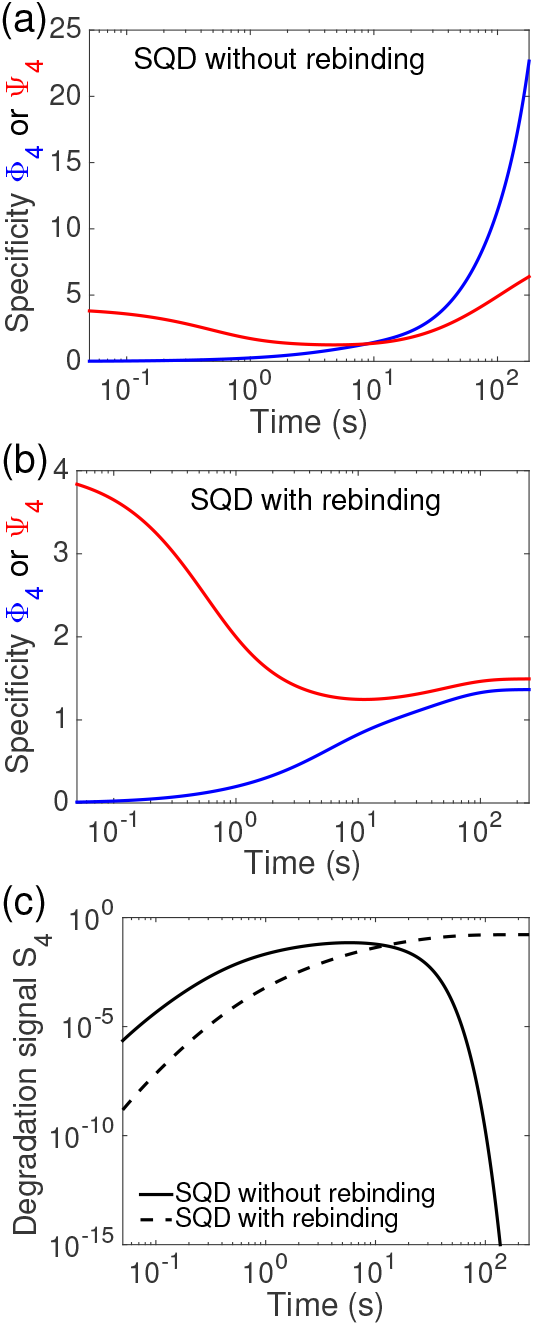
Time dependence of specificities in SQD model with and without reversible binding to E3. Time trajectories of Φ_4_ (blue) and Ψ_4_ (red) generated by (a) the SQD model without rebinding (i.e., substrate-E3 single-contact model in Ref. [10]. Initial condition: *c*_0_ = 1) and (b) the SQD model with rebinding (initial condition: *s*_0_ = 1). (c) Time trajectories of degradation signal *S*_4_ (simulations started from initial state *S*_0_). To be consistent with the analysis in Ref. [10], Φ_4_ was computed as Φ_4_ = -*∂* log(*S*_4_ + *C*_4_)/*∂* log *k* (similarly to Ψ_4_). *k_b_* = 0.5 s^-1^, [E3]=8 nM, and *k*_on_ = 0.01 nM^-1^s^-1^. Beyond 200 seconds, the degradation signal in the model without rebinding is too small to be quantified with sufficient numerical accuracy, and consequently the computation of specificities becomes numerically unstable.

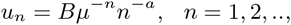

where *u_n_* is the elongation rate from state *c_n_* to state *c*_*n*+1_. To resolve parameters *B, μ* and *α*, we fit the chain length distributions for both SCF substrates cyclin E and *β*-catenin reported in [13] with a fixed k. Without E3-bound DUB editing (complying with the experiment in Ref. [13]), the chain length distribution can be calculated

**FIG. S7.**
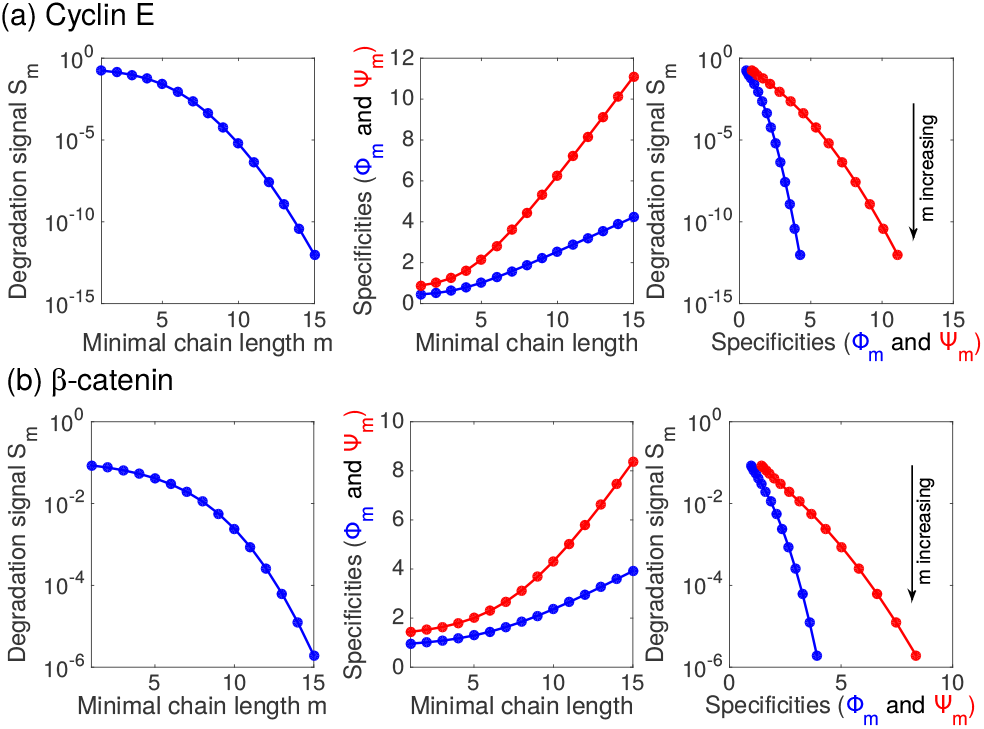
The minimal signaling chain length. Degradation signal *S_m_*, specificities Φ_*m*_ and Ψ_*m*_ and their tradeoffs as functions of the minimal chain length m, in SCF substrates (a) Cyclin E and (b) *β*-catenin. The SQD model with E3-bound DUB editing (*η* = 1) were simulated with *k_b_* = 0.5 s^-1^ and [E3]=8 nM. The ubiquitin transfer rates *k_u_*’s are provided in the caption of Fig. 4.

as:

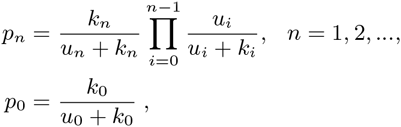

where *p_n_* is the probability for a substrate molecule to conjugate a chain of length *n. k_n_* = *k, n* = 0, 1,….

The SAW model can be potentially used to explain ligase interacting with more than one E2s. For the ligase APC, ubiquitin chain initiation is mediated by E2 UbcH10 that can rapidly conjugate on average 2-4 ubiquitins to a substrate [3], followed by a slower elongation mediated by UbeS2. This kinetic difference between UbcH10 and Ube2S might be caused by their differences in structure and distance to the acceptor sites. Especially in the SAW model, the number of reactive chain configurations depends on the distance between the E2 and substrate on the ligase. UbcH10 and Ube2S interact with APC simultaneously via different binding domains (therefore locate at difference distances from the substrate) and could independently (but complement to each other) function with a slow initiation and an elongation slowdown, only by SAW models with difference parameter values. UbcH10 has a faster early elongation rate which steeply declines after a smaller number of ubiquitins being transferred to the substrate. In comparison, Ube2S might have a slower elongation with a more gradual length-dependent slowdown, allowing elongation of a longer chain.

### Temporal dynamics of degradation and specificities

The model in Ref. [10] compared transient ubiquitination dynamics of CD4-M-Vpu and CD4-Vpu pairs after a single substrate-E3 contact, without considering the substrate rebinding to the E3. It is worth noting that comparing polyubiquitination levels in wild-type CD4 and CD4-M by this model is less robust as demonstrated in Fig. S6a that Φ_4_ increases unboundedly as time evolves and the polyubiquitination signal *S*_4_ + *C*_4_ continuously declines, making it impossible to quantitatively assess substrate discrimination by a single time point. By contrast, values of Φ_4_ from temporal samples generated by the SQD model (*η* = 1) are much smaller at corresponding time points. Modified substrate binding back to E3 ligase in the SQD model allows the substrate to re-enter the ubiquitin chain elongation and eventually stabilize the signal level. Varied DUB rates in the SQD model generated diverse dynamics of Φ_4_, and the DUB dose response could even differ in different time points (Fig. S6(d)). At low DUB activity (*k_b_* = 0.05 s^-1^), substrate discrimination is only visible during the transient dynamics, reaches a maximum at an intermediate time (i.e., the signal is most sensitive to perturbations in *k*), and diminishes in the steady state when all substrates are elongated beyond the signaling state.

**FIG. S8.**
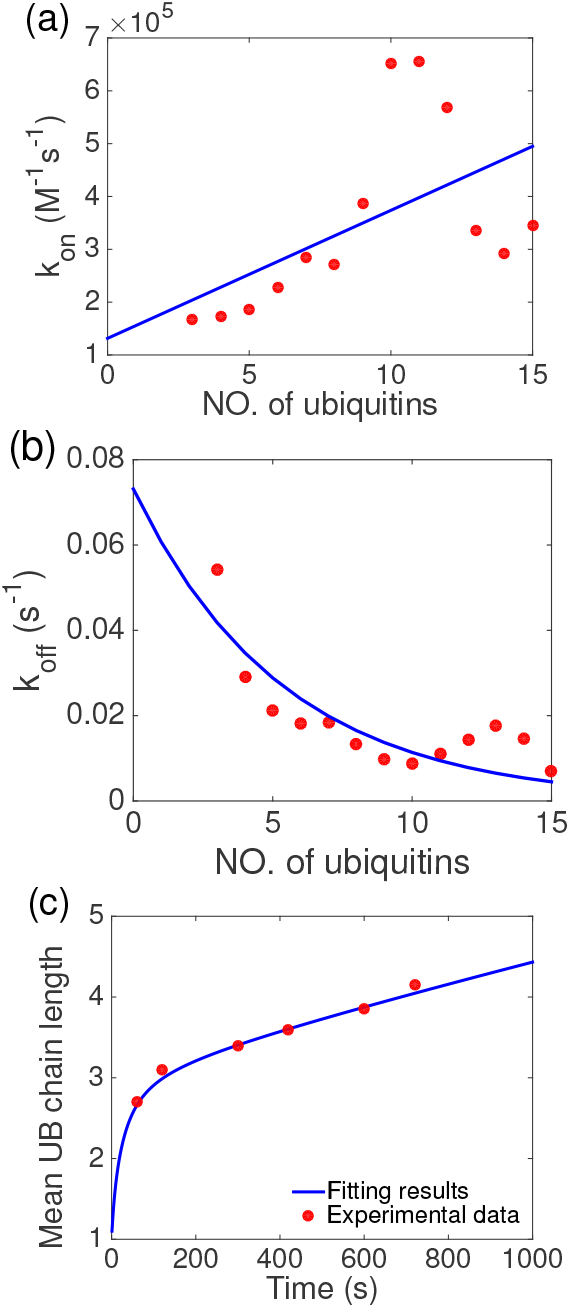
Least-square fits to APC-securin association, dissociation rate constants, and mean chain length dynamics data from Lu *et al.* [3]. Chain length-dependent association and dissociation rate constants of substrate securin binding to ligase APC were measured by the single-molecule fluorescence experiments. (a) The association rate constant fits the data by a linear function of the chain length: *k*_on,*n*_ = (0.24*n* + 1.3) x 10^−4^ nM^-1^s^-1^. (b) The dissociation rate constant fits to data with an exponential function of the chain length: *k*_off,*n*_ = 0.073*e*^-0-186*n*^ *s*^-1^. (c) Fitting to the time trajectory of the mean chain length from the single-contact assay to obtain the elongation rates: *k_u,n_* = 0.38 x 1.2^-n^ x *n*^-0.92^ *s*^-1^, by the assumption *k*_*u*,0_ = 0.05*k*_*u*,1_.

### Stage-dependence of E3-substrate binding – Processive affinity amplification

The stage-dependence of substrate affinity to the E3 ligase was observed by Lu *et al.* [3] using single-molecule fluorescent assays. We use a linear function to fit the dependence of association rate constant *k*_on,*n*_ of securin to APC on the chain length (Fig. S8a. Note that noise in the data is visible when n is large, possibly due to small number of data points). The dissociation rate constant *k*_off,*n*_ is fit to a decaying exponential function of the chain length (Fig. S8b). temporal dynamics of the mean chain length is fit by simulating an ODE model of SQD under the conditions of slow chain initiation and elongation slowdown by the SAW model (Fig. S8c). We note that the information about the chain initiation rate *k*_*u*,0_ is not available from Ref. [3] and we therefore assume that the initiation rate is 20 fold smaller than the rate of the next elongation step. It might be possible to resolve *k*_*u*,0_ from the single-molecule kinetics provided enough trajectory data. With a known *k*_off,0_, *k*_*u*,0_ can be calculated from the average number of substrate-E3 encounters before a successful conjugation *n_c_* = (*k*_off,0_ + *k*_*u*,0_)/*k*_*u*,0_ (or from the fraction of encounters that produces the first transfer [13]).

**FIG. S9.**
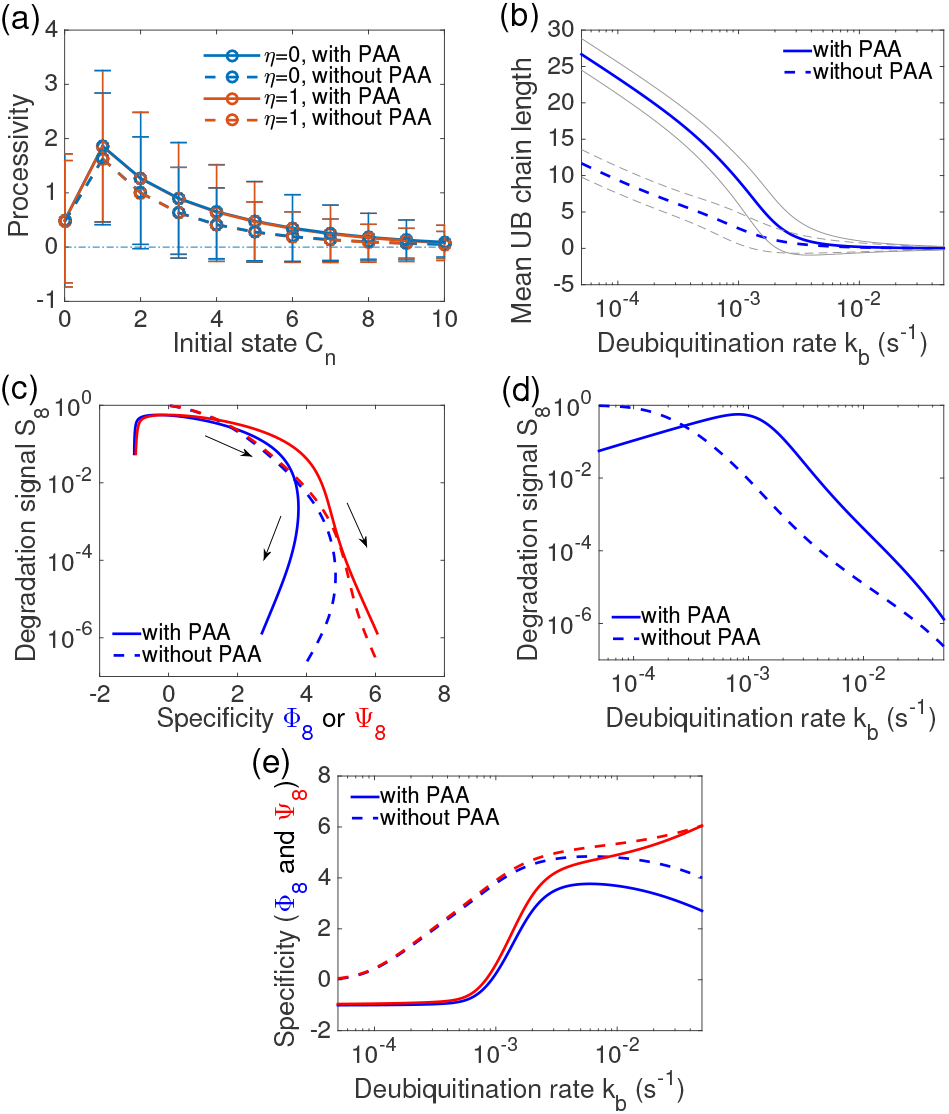
Effects of processive affinity amplification on degradation signal and specificities, at the minimal signaling chain length *m* = 8. (a) Stage-dependent processivity (*k_b_* = 5 x 10^−4^ s^-1^), in comparison to results by kb = 0.02 s^-1^ shown in Fig. 6a. (b) The mean chain length 〈*n*〉_*s*_. The standard deviations are shown as grey outlines. (c) Signal-specificity tradeoffs. Points at *k_b_* = 5 x 10^−4^ s^-1^ are marked by ‘x’. Arrows indicate the increasing direction of *k_b_*. (d) Signal *S*_8_ and (e) specificities Φ_8_ and Ψ_8_ as *k_b_* varies. Equations that fit *k*_off,*n*_, *k*_on,*n*_, and *k*_*u,n*_ are shown in Fig. S8. Except for panel (a), all results were obtained from the SQD model with E3-bound DUB editing. For the null model without PAA, stage-independent *k_on_* = *k*_*on*,0_ = 0.00013 nM^-1^s^-1^ and *k*_off_ = 0.073 s^-1^ were used. [E3] = 10 nM.

Fig. S9 shows results under the minimal signaling chain length *m* = 8. The processivity differences between models with or without PAA (*η* = 0 or 1) under a small DUB rate *k_b_* = 5 x 10^−4^ s^-1^ (compared to *k_b_* = 0.02 s^-1^ in Fig. 6a in the main text) are almost indiscernible (Fig. S9a). In the case of PAA, the system achieves a maximum mean chain length 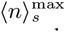 at an optimal DUB activity, however the PAA also produces much bigger fluctuations in chain length across a wide range of DUB activity (Fig. S9b). At low DUB activity, even though most substrate molecules are polyubiquitinated, the PAA causes higher substrate buffering by the E3 ligase and therefore produces lower signal than the case with no PAA (Fig. S9b and d). High DUB activity inhibits ubiquitination as expected. Specificity switches are more pronounced under the PAA (Fig. 6e)

**TABLE S3.** Specificities in model variants for the SCF substrates: *β*-catenin and Cyclin E1

| | Model | $S_4$ | $\Phi_4$ | $R_{0.5}$ | $R_{10}$ | $\Psi_4$ | $R_{0.5}$ | $R_{10}$ | $\Phi_1$ | $\Psi_1$ |
| --- | --- | --- | --- | --- | --- | --- | --- | --- | --- | --- |
| β-cat | EBD ($\eta=0$) | $9.7 \times 10^{-3}$ | 0.83 | 1.45 | 21.7 | 1.05 | 2.19 | 26.8 | 0.69 | 0.90 |
| | EBD ($\eta=1$) | $2.7 \times 10^{-3}$ | 0.25 | 1.16 | 8.7 | 1.36 | 2.83 | 94.3 | 0.11 | 0.96 |
| | SQD ($\eta=0$) | $8.1 \times 10^{-2}$ | 1.30 | 1.78 | 77.4 | 1.51 | 3.16 | 95.5 | 1.07 | 1.28 |
| | SQD ($\eta=1$) | $5.4 \times 10^{-2}$ | 1.17 | 1.69 | 62.3 | 1.79 | 4.10 | 492 | 0.96 | 1.44 |
| CycE | EBD ($\eta=0$) | $3.4 \times 10^{-2}$ | 0.66 | 1.38 | 30.8 | 0.87 | 2.09 | 38.0 | 0.35 | 0.56 |
| | EBD ($\eta=1$) | $9.6 \times 10^{-3}$ | 0.30 | 1.19 | 13.3 | 1.51 | 3.35 | 203 | 0.06 | 0.79 |
| | SQD ($\eta=0$) | $1.1 \times 10^{-1}$ | 0.94 | 1.56 | 62.7 | 1.15 | 2.59 | 73.3 | 0.53 | 0.74 |
| | SQD ($\eta=1$) | $5.6 \times 10^{-2}$ | 0.79 | 1.47 | 45.0 | 1.61 | 3.89 | 531 | 0.45 | 0.89 |

– We note that $\Phi_4$ and $\Psi_4$ are more accurate estimates to perturbations by small parameter changes (better approximations to $R_{0.5}$ than to $R_{10}$ even though their trends are identical).

*en bloc* **ubiquitin transfer**. Here we use the SQD model without E3-bound DUB editing (*η* = 0) to briefly discuss the scenario of *en bloc* ubiquitin transfer under uniform process rates. Assume on average a ligase system conjugates *z* ubiquitins to a substrate per transfer event. The processivity is now *ρ* = *zu/k*. Given the minimal signaling chain length m, the minimal transfer steps required to reach signaling states is *r* = [*m/z*], which is the minimal integer equal to or greater than *m/z* (e.g., if *z* = 3 and *m* = 4, *r* = 2 is the minimal number of steps to the signaling state). For simplicity, we assume the deubiquitination also removes *z* ubiquitin molecules from a substrate at a time. The backward processivity for the dissociated substrate is *ρ_b_* = *zb*. Following the development in the previous section, we can write general equations of degradation signal and specificities in terms of processivity *ρ* and *ρ_b_* (under the condition of bounded elongation, *ρ* < *ρb*):

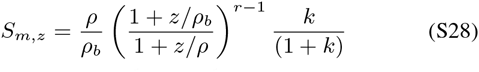

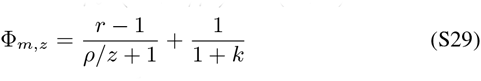

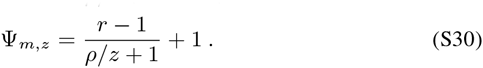

The sequential transfer model is recovered when *z* = 1. Assume the processivity *ρ* and *ρ_b_* remain relatively unchanged for different *z*. Under this circumstance, one can verify that *en bloc* transfer (increasing *z*) reduces specificities and in the meantime less intuitively also decreases the degradation signal.

### Numerical computation of model quantities

Models that do not have analytical results rely on numerical simulation, which include the parsimonious SQD model when *η* = 1 and all models parameterized with nonuniform rates. The steady-state quantities of a model can be obtained given the steady-state probability distribution over states *s_n_* and *c_n_, n* = 0,1…,. Numerical integration of the master equations to the steady state is a straight forward computation, which however requires to determine whether a simulation reaches its steady state (not always numerically robust). By an alternative, we compute the stationary probability distribution by seeking the null vector of the generator matrix of the master equation for a model with a finite number of states. Consider a model of *N*-stage ubiquitination, i.e., with states *s_n_* and *c_n_, n* = 0,1,…, *N* – 1. The model has 2*N* states in total. The master equation can be written as:

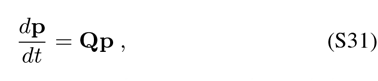

where **p** is a 2*N* x 1 column vector that contains the probability distribution over states at time *t*. The 2*N*-by-2*N* matrix **Q** is the generator matrix that contains transition rates between states. For an EBD or a SQD model, **Q** has a one dimensional null space and the steady-state solution to 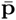 is a right-side null vector. The degradation signal *S_m_* can be obtained directly from 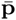. In practice, the distribution **p** and the generator matrix **Q** matrix can be arranged in a specific order for analytical and computational convenience. Let **p** = [*s*_0_, *s*_1_,…, *s*_*N*-1_, *c*_0_, *c*_1_, *c*_*N*-1_]^*T*^. The **Q** matrix can then be partitioned into 4 smaller square matrices:

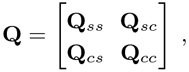

where the subscript indicates whether the submatrix contains rate constants of transitions within unbounded or E3-bounded states or between unbounded and E3-bounded states. **Q**_*cs*_, for example, contains rate constants of transitions from states *s_i_,i* = 0,1,…,*N* – 1 to states *c_j_, j* = 0,1,…, *N* – 1. Under uniform and normalized rates, these matrices in the SQD model at *N* = 4, for example, are constructed as:

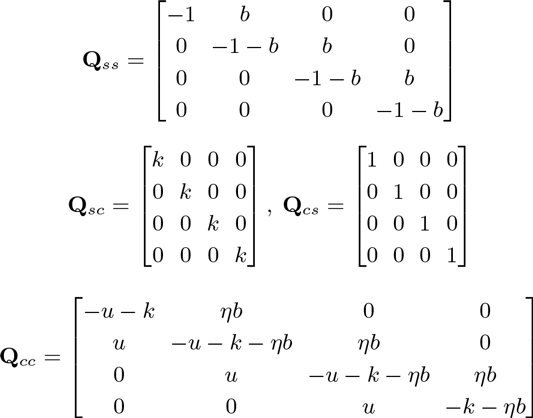

Given the conservation of probability (i.e., all entries in **p** sum up to 1), 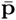 can be solved by replacing an arbitrary row (say the rth row) of **Q** with a row of 1’s to form a new matrix **Q**_*r*_ equation:

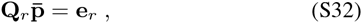

where e_*r*_ is the elementary vector with 1 at the *r*th entry. Therefore, 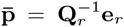. The degradation signal *S_m_* is a partial sum of the vector 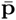. A solution in general depends on the system size *N* and asymptotically converges when N becomes large.

Specificities Φ_*m*_ and Ψ_*m*_ can be obtained as follow. Taking partial derivative over both sides of Eq. S31 with respect to *k*, we have

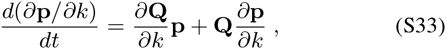

with a constraint Σ(*∂***p**_*i*_/*∂k*) = 0. Once 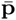 has been obtained, the steady-state specificity Φ_*m*_ can be calculated by solving for *∂***p**/*∂k* = -**Q**_*r*_**b**, where **b** is the vector 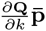 with the *r*th entry replaced with 0. Thus, we have

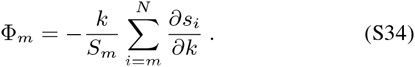

The specificity Ψ_*m*_ can be similarly obtained. Dynamics of specificities Φ_*m*_ (*t*) and Ψ_*m*_ (*t*) can be solved by ODE integration of Eq. S33 together with the master equation Eq. S31 given proper initiation conditions.

When ubiquitin transfer or substrate dissociation has nonuniform rates, Φ_m_ and Ψ_*m*_ are calculated as relative sensitivity of a scaling factor *w* of all rates at *w* = 1. For the case of nonuniform ubiquitin transfer rates:

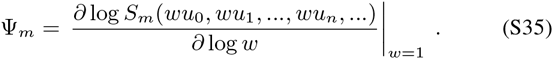

The generator matrix block *Q_cc_* is formed as:

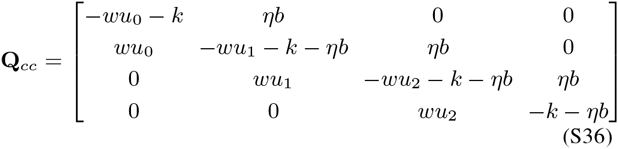

The Eq. S33 is rewritten as

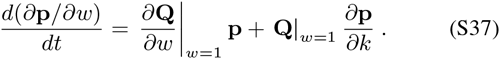

For the case of nonuniform dissociation rate constants:

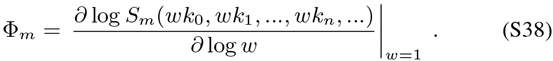

